# Engineering prolyl hydroxylase-dependent proteolysis enables the orthogonal control of hypoxia responses in plants

**DOI:** 10.1101/2024.12.13.628401

**Authors:** Vinay Shukla, Sergio Iacopino, Laura Dalle Carbonare, Yuming He, Alessia Del Chiaro, Antonis Papachristodoulou, Beatrice Giuntoli, Francesco Licausi

## Abstract

Vascular plants and metazoans use selective proteolysis of transcription factors to control the adaptive responses to hypoxia, although through distinct biochemical mechanisms. The reason for this divergence is puzzling, especially when considering that the molecular components necessary to establish both strategies are conserved across the two kingdoms. To explore an alternative evolutionary scenario where plants sense hypoxia as animals do, we engineered a three-components system aimed to target proteins for degradation in an oxygen dependent manner *in Arabidopsis thaliana*. Applying the synthetic biology framework, we produced a hypoxia-responsive switch independent of endogenous pathways. When applied to control transcription, the synthetic system partially restored hypoxia responsiveness in oxygen-insensitive mutants. Additionally, we demonstrated its potential to regulate growth under flood-induced hypoxia. Our work highlights the use of synthetic biology to reprogram signalling pathways in plants, providing insights into the evolution of oxygen sensing and offering tools for crop improvement under stress conditions.

## Introduction

Aerobic organisms require molecular oxygen (O_2_) for sufficient ATP synthesis to support growth and development. When O_2_ provision to cells from the surrounding environment is limited (hypoxia), cells adjust their structure, physiology and metabolism to avoid or resist the stress. Such adaptations require transcriptional reprogramming triggered by dedicated hypoxia sensing mechanisms. Remarkably, vascular plants and metazoans share a similar sensing strategy based on oxygen-dependent degradation of constitutively expressed transcription factors (TFs) and their nuclear accumulation upon exposure to hypoxia^1^. However, this is achieved by different biochemical solutions in the two kingdoms.

In vascular plants, a major role in transcriptional reprogramming is played by the Ethylene Response Factor VIIs (ERFVIIs), TFs controlled via the Plant Cysteine Oxidase (PCO)-branch of the N-degron pathway^2–4^. This is a proteasomal degradation pathway that dictates protein stability depending on the exposed N-terminal amino acid^5^. Methionine amino peptidases prepare the ERFVIIs to expose an N-terminal cysteinyl residue for N-terminal sulfination via PCOs^6^. This modification promotes N-terminal arginylation, which is in turn recognised by the ubiquitin E3 ligase PROTEOLYSIS6 (PRT6), with the assistance of BIG, for polyubiquitination and the consequent degradation through the proteasome^7–9^ (**Supplementary Fig. 1a)**.

In mammalian cells, transcriptional adaptation to low O_2_ conditions is mediated by the Hypoxia-Inducible Factors (HIFs), which belong to the basic helix-loop-helix protein family^10^. They are dimers of alpha and beta subunits, the former controlled via oxygen-dependent degradation^11,12^. This regulation is mediated by the hydroxylation of internal prolyl residues (Pro402 and Pro564 in HIF-1α), catalysed by O_2_ and 2-oxoglutarate (2-OG)-dependent enzymes Prolyl-hydroxylases (PHDs)^13,14^. The hydroxylated HIFα subunit is recognised by a ubiquitin E3 ligase complex that contains the substrate recognition subunit von Hippel Lindau factor (VHL)^15,16^. As in the case of ERFVIIs, polyubiquitination leads to HIFα degradation through the proteasome complex. When oxygen levels drop, instead, HIFα is protected from proteolysis and thereby accumulates in the nucleus. Here, heterodimerization with its cognate β-subunit reconstitutes a functional transcription complex able to induce expression of hypoxic genes (**Supplementary Fig. 1b)**.

The evident similarity between vascular plant strategy and metazoan O_2_ sensing strategy has led to the speculation that this represents the best solution to accommodate organised multicellularity^17^. However, the last common ancestor of plants and animals likely expressed both PHD and N-Cys dioxygenase^18^. This raises the question as to whether PHD adoption better suits O_2_ sensing in heterotrophs with an active system for gas circulation, whereas N-Cys dioxygenases optimally accommodate photosynthetic organisms. The recruitment of oxygenases with distinct kinetic parameters as O_2_ sensors could reflect substantial differences in the responses required in hypoxia in the two kingdoms, in the O_2_ levels at which these need to be activated and the existence of peculiar mechanisms of interference or crosstalk with metabolites and secondary messengers^19–21^.

Hypotheses regarding the origin and differentiation of molecular mechanisms are traditionally addressed by comparisons across extant species. We reasoned that a synthetic biology approach would instead tackle more directly the question about the convergence/divergence of O_2_ sensing mechanisms in animals and plants ^22^. Therefore, we set out to reconstruct a hypoxia responsive system based on proteolysis for plant cells based on proline hydroxylation. Previously, we exploited the O_2_-dependent interaction of HIF and VHL to engineer a molecular circuit able to inhibit gene expression in hypoxia^23^. Here, we re-engineered O_2_-dependent proteolysis to control the stability of reporter and signal transducers. Besides addressing the fundamental question regarding the evolution and diversification of O_2_ sensing in multicellular eukaryotes, we considered that such an effort would serve as a proof of concept for the engineering of proteostatic signalling in plant cells^24^. Moreover, by doing so, we thought to establish an orthogonal switch to control the response to environmental conditions that limit O_2_ availability for plants, such as submergence.

Flooding is one the main causes for agricultural losses worldwide^25^. Since gas diffusion is severely reduced in water, submerged plant organs experience limited O_2_ availability. Tinkering/tampering with the endogenous oxygen sensing system is likely to compromise plant fitness overall. This is because the N-cys degron pathway controls the response to different environmental stresses including cold, high salinity, pathogen attack and dehydration^26–28^ and also participates to developmental processes^29–32^. We therefore applied the synthetic biology framework to generate an orthogonal switch for transcription regulation in response to hypoxia in plant cells. We used this molecular device to explore the alternative evolutive scenarios where O_2_ sensing in plants depends on PHDs and tested its ability to drive developmental responses to submergence-induced hypoxia.

## Results

### Engineering PHD-dependent proteolysis in plant cells

We set out to reconstruct an orthogonal O_2_-dependent molecular switch for plant cells by inducing targeted proteolysis based on the mammalian HIF-1α/VHL system (**Supplementary Fig. 1B**). We have shown previously that the C-terminal Oxygen-dependent degron of HIF-1α (HIF_ODD_), and the VHL’s β-domain (aa63-157) interact in an O_2_-dependent manner exclusively in the presence of a human PHD3 enzyme in *Arabidopsis thaliana* cells^23^. Given the 1.7-1.4 billion years evolutionary distance between plants and animals^33^, we considered unlikely for the mammalian VHL to associate with functional E3 ligase complexes in Arabidopsis cells. An initial comparison of *in silico* reconstructed Cullin-Ring E3 ligase (CRLs) complexes from animals and plants suggested similar subunit arrangements between the complex responsible for HIF-α hydroxylation in human cells and those dedicated to auxin signalling in Arabidopsis (**Fig. 1A**). However, while VHL contacts the CRL complex via the interaction between its α-domain with ElonginB (ELOB) and ELOC ^34^ (**Fig. 1A**), the auxin sensor Transport Inhibitor Response1 (TIR1) does so by binding to the Arabidopsis homolog of the mammalian S-phase kinase associated protein 1 (Skp1), Ask1 ^35^ (**Fig. 1A**). Moreover, the α-domain of VHL resembles TIR1 F-box, with a similarly oriented three-helical structure; nonetheless, VHL’s inability to contact Ask1 in Arabidopsis cells is supported by the distinct amino acid residues involved in VHL/ELOC and F-box/Ask1 interaction, coupled with other minor structural differences (**Supplementary Figure 2A and 2B**), and with the absence of a Cullin2 homolog in plants^36^ . Based on these observations, we decided to replace the original VHL α-domain with an endogenous plant F-box domain. Under aerobic conditions, the chimeric F-box protein is expected to bind the hydroxylated HIF_ODD_ peptide, thereby promoting the ubiquitination and degradation of the fusion protein in which it is contained. Under hypoxia, instead, binding of VHL to the HIF_ODD_ peptide should be prevented, thus stabilising the HIF _ODD_ -tagged protein (**Fig. 1B**).

**Fig. 1:**
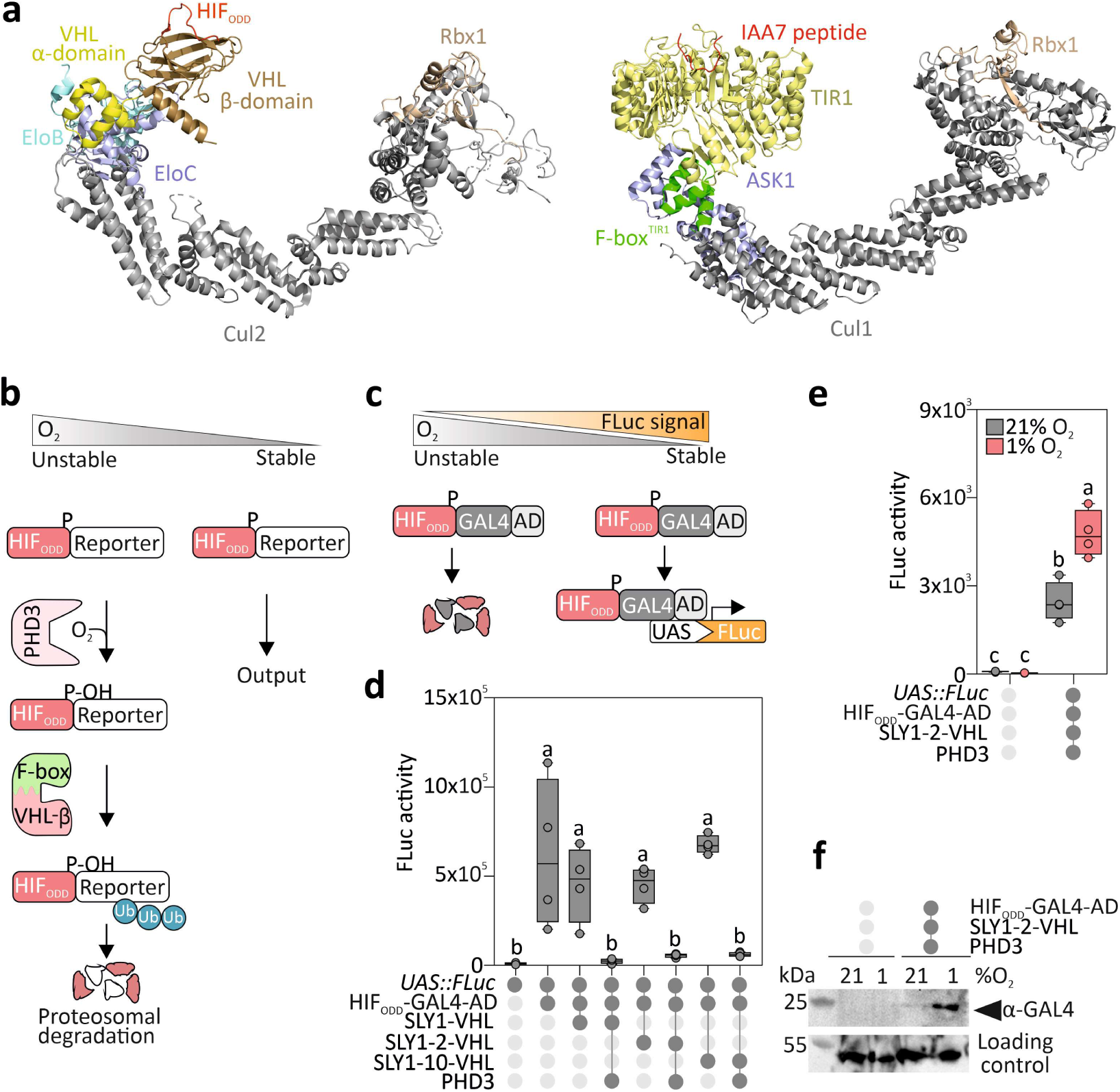
Implementation of PHD- and VHL-dependent O_2-_sensing in plant cells. **a,** Comparison of the predicted human VBC-CR (top) and plants SCF^Tir1^ (bottom) complexes. The VBC-CR complex comprises the scaffold protein Cul2 (grey), the E3 ubiquitin ligase Rbx1 (light brown), the adaptor proteins EloB (aquamarine) and EloC (blue) and the substrate recognition protein VHL (α-domain in yellow, β-domain in bronze). The plant SCF complex consists of the Cul1 scaffold protein (grey), Rbx1 (light brown), the adaptor Ask1 (blue) and the substrate recognition F-box protein TIR1 (LRR region in yellow, F-box domain in green). The peptides shown for the animal and plant substrates are HIF_ODD_ and IAA7, respectively. **b,** The O_2_-dependent proteolysis mechanism designed for plant cells. A chimeric E3 ligase, which comprises the VHL beta domain and a plant F-box domain, recognizes the HIF_ODD_ domain when this is hydroxylated by PHD in the presence of O_2_. Reporter proteins linked to the HIF-ODD are degraded in normoxia but stable in hypoxia. **c,** Reporter strategy deployed to test the synthetic O_2_ sensing system. HIF_ODD_ was fused to the DNA binding domain of GAL4 and the activation domain (AD) of RAP2.12. In the presence of O_2_, this chimeric transcription factor is degraded by the 26S proteasome. In hypoxic conditions, instead, it is stabilised and binds the *4xUAS* promoter, to activate transcription of a firefly luciferase reporter (FLuc). **d,** FLuc output measured in Arabidopsis mesophyll protoplasts transformed with combinations of plasmids to express HIF_ODD_-GAL4-AD, three variants of the SLY1-VHL chimera (**Supplementary Fig. 4a**) and PHD3. **e,** Effect of hypoxia (1% O_2_, 6 h) on FLuc output in protoplasts expressing HIF_ODD_-GAL4-AD, SLY1-2-VHL and PHD3. **f,** Immunodetection of HIF_ODD_-GAL4-AD in the same experimental setup as **e**. Letters indicate statistically significant differences (p ≤ 0.05), as determined by one-way ANOVA followed by Tukey’s post-hoc test (n = 4).

We first attempted to generate the desired chimeric F-box protein by fusing the TIR1 F-box domain to the VHL β-domain. To monitor HIF degradation, we produced a chimeric transcription factor consisting of the HIF_ODD_ peptide, the minimal GAL4 DNA binding domain and the small but potent activation domain of the Arabidopsis transcription factor Related to APETALA2.12 (RAP2.12) ^37^ (**Fig. 1C**). This construct was expected to bind the synthetic 4xUAS promoter^23^ and promote the transcription of a Firefly Luciferase (FLuc) reporter gene (**Fig. 1C**). We have shown previously that, in Arabidopsis cells, hydroxylation of the HIF_ODD_ peptide requires expression of a mammalian PHD enzyme^23^. Therefore, we speculated that, in the presence of O_2_ and PHD3 from *Homo sapiens*, the HIF transcription factor can be hydroxylated and recognised by the chimeric VHL RCL complex for ubiquitination and proteasomal degradation (**Fig. 1C**). When expressed in Arabidopsis protoplasts, HIF-GAL4-RAPAD induced FLuc reporter activity, but this was not significantly affected by co-expression of the other VHL-F-box and PHD3 modules (**Supplementary Fig. 3A-B**). This result suggested the inability of the TIR1-based chimeric F-box protein to contact the HIF_ODD_ peptide, independently from its hydroxylation state. Therefore, we tested a second F-box domain from the SLEEPY1 (SLY1) protein, which participates in gibberellin (GA) signalling^38^. We chose SLY1 because it is one of the smallest functional F-box proteins in the Arabidopsis proteome ^39^. In addition to this, two gibberellin insensitive SLY1 variants, SLY1-2 and SLY1-10, have been characterised as early protein truncations that abolish gibberellin responsiveness while maintaining intact the F-box domain^38^ (**Supplementary Fig. 4A**). Since no crystal structure for SLY1 is available, we superimposed the predicted model of its F-box domain on TIR1 to verify the conserved identity and position of residues necessary for the interaction with Ask1 (**Supplementary Fig. 4B**). We fused the wild type and truncated SLY1 variants at the N-terminal end of the VHL β-domain and used them in a protoplast transactivation assay, as described before for TIR1. This time, combination of SLY1-VHL and PHD3 effectors abolished the ability of HIF_ODD_-GAL4-AD to enhance reporter activity, indicating SLY1-VHL their efficacy at ubiquitinating the hydroxylated HIF-GAL4-AD transcription factor (**Fig. 1D**). All SLY1 versions, including the two truncated fragments, were able to repress Fluc expression. We thus selected the smallest fragment, SLY1-2, for further experiments, under the assumption that this mutant will likely interfere the least with GA signalling. An N-terminal GFP fusion to SLY1-2-VHL showed co-localization of GFP and DAPI signals in Arabidopsis protoplasts (**Supplementary Fig. 4C**). This validated the predicted nuclear localisation of SLY1-2-VHL. and confirmed its potential to recognise the HIF-GAL4-AD substrate in the nucleus. We then used the selected combination of effectors to test whether treatment of protoplasts with hypoxia (0.1% O_2_ for 6 hours) could alleviate the repression imposed by SLY1-2-VHL on HIF_ODD_-GAL4-AD activity and observed a significant increase of reporter output under hypoxia (**Fig. 1E**). This suggests that the degradation of HIF_ODD_-GAL4-AD is inhibited when PHD3 activity is limited by insufficient co-substrate availability. This hypothesis was corroborated by immunodetection of GAL4-AD in a western blot (**Fig. 1F**).

### HD-dependent conditional proteolysis is transferable to different reporter proteins

We studied whether the newly established O_2_-dependent proteolytic mechanism can be generally used to control the stability of other proteins to generate hypoxia reporters. To this end, we generated single transcriptional units coding for two luciferases (defined as Luc1 and Luc2, for upstream and downstream) separated by a Ubiquitin10 monomer. We fused the HIF_ODD_ domain to the Luc2 so that, during mRNA translation, deubiquitinases (DUBs) would release two fragments: a stable Luc1-UBQ fragment, suitable for signal normalization, and a HIF-fused Luc2 proteoform, whose abundance is dependent on O_2_ and PHD3 expression (**Supplementary Fig. 5A**). We set out to test variants of this construct, each differing for a combination of three features: the 5’ untranslated mRNA region (UTR), the paired luciferases and the number of HIF_ODD_ motifs associated with Luc2. The different reporter variants were evaluated in a protoplast testbed. For every variant, we generated two independent transgenic lines of Arabidopsis stably expressing the SLY1-2-VHL effector along with the reporter. Mesophyll protoplasts isolated from each line were then transformed with a *35S:PHD3* construct or *35S:GFP* as a control (**Supplementary Fig. 5B**). We measured the relative luciferase signal ratio produced in protoplasts after 12 h incubation in normoxic (21% O_2_) or hypoxic (1% O_2_) conditions, screening for reporter versions characterized by decreased output in case of PHD3 expression under normoxia (as compared with their normoxic output in the absence of this effector), and high output recovery under hypoxia. As the best performing reporter of our set, we thereby identified, the construct consisting of a *Renilla reniformis* luciferase (RLuc) as a normaliser and FLuc as a reporter, driven by a 35S CaMV promoter and equipped with a single HIF_ODD_ at the C-terminal end (**Fig. 2A-B**). We named this O_2_-dependent ratiometric oxygen sensor as O_2_ratio. All other variants proved to have little responsiveness to either PHD3 or hypoxia (**Supplementary Fig. 5C**). We also tested whether we could increase the dynamic range of this reporter between normoxia and hypoxia by rational mutagenesis of PHD, aimed at lowering its activity under O_2_ limitations. Inspired by past studies on the human PHD2 enzyme^40–42^, we tested three different single amino-acid substitutions (**Supplementary Fig. 6A-B**). In protoplasts transfected with the same strategy as above, the wild type PHD3 showed the best dynamic range among the four variants, with consistently significant differences between normoxia and hypoxia in both independent *SLY1-2-VHL/O_2_ratio* lines (**Supplementary Fig. 6C-D**). We then proceeded to transform one of the two *SLY1-2-VHL/O_2_ratio* lines with a stable *35S:PHD3* construct (or its mutant versions, **Supplementary Fig. 6E**) and tested two independent lines for each. We confirmed that expression of the wild type PHD3 was associated with the highest range of output reduction in normoxia and a significant output recovery after 12 h hypoxia in comparison with normoxic controls (**Fig. 2B, Supplementary Fig. 6F**). Neither of the two *35S:PHD3* lines showed developmental or growth differences in comparison with wild type plants, indicating substantial orthogonality of the system (**Fig. 2C**). Looking at O2ratio dynamics in *PHD3*-expressing plants under normoxia and hypoxia, we found that the earliest significant output increase occurred after 6 h hypoxia (**Fig. 2D**).

**Fig. 2:**
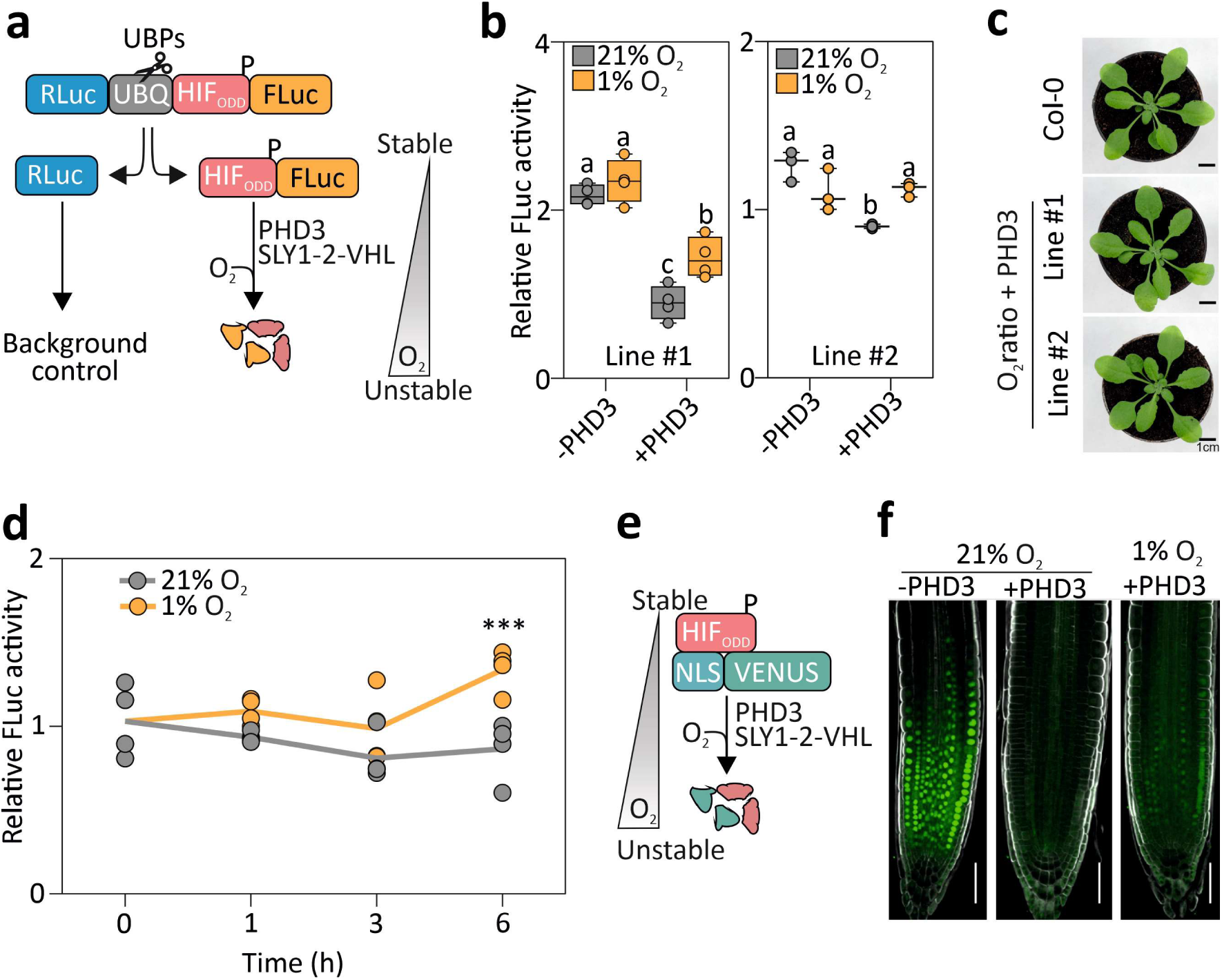
Design and testing of a ratiometric PHD- and VHL-dependent O₂ reporter in plants. **a,** O_2_ratio, a polyprotein consisting of Renilla Luciferase (RLuc), a UBQ10 unit and FLuc-HIF_ODD_, was expressed in plant cells together with PHD3 and SLY1-2-VHL (not shown). Endogenous ubiquitin proteases (UBPs) co-translationally cleave the polypetide at the c-terminus of UBQ10. The resulting Rluc serves as a normalization reference while FLuc-HIF_ODD_ is used to monitor O_2_ levels. **b,** PHD- and O_2_-dependent stability of O_2_ratio in plants that express the modules described in A. Letters indicate statistical differences (p ≤ 0.05) determined by two-way ANOVA followed by Tukey’s post-hoc test (n = 4). **c,** Phenotype of representative 4-week-old O_2_ratio and wild-type Col-0 plants (scale – 1cm). **d,** O_2r_atio output dynamics during a hypoxia (1% O_2_, yellow) or normoxia (21% O_2_, gray) time course. The relative luminescence output was normalized to the average FLuc/RLuc value at time 0 h. Letters indicate statistical difference (p≤0.05) between treatments at each time point, as assessed by one-way ANOVA followed by Tukey’s post-test (n=4). **e,** The fluorescent protein Venus, fused with a nuclear localization signal (NLS) and HIF_ODD_, serves as a reporter of O₂ levels. **f,** Stability of the Venus fluorescence signal when regulated by PHD activity and atmospheric O₂ availability in root tips of 7d old Arabidopsis seedlings subjected to 21% or 1% O_2_ for 6 hours (scale – 50 µm).

Finally, we tested the portability of the regulation by transferring the HIF_ODD_ degron to a nucleus-targeted VENUS reporter, expressed under the control of the p16 promoter (**Fig. 2E**)^43^. Using confocal microscopy, we could detect a strong fluorescent signal in the root tip of transgenic plants expressing this reporter construct, which disappeared almost completely after expression of a *PHD3* transgene (**Fig. 2F**). This latter transgenic line recovered nuclear fluorescence when plants were incubated for 6 h at 1% O_2_ (**Fig. 2F**). We concluded that the combination of three modules of human origin (*PHD3*, *SLY1-2-VHL* and *HIF_ODD_*) is a broadly transferable system to control the stability of reporter proteins in plants depending on O_2_-availability.

### PHD-dependent proteolysis can control transcriptional responses to hypoxia in plant cells

The possibility to establish O_2_-dependent, targeted proteolysis allowed us to test an evolutionary scenario where hypoxia signalling did not diverge between vascular plants and metazoans. We did so by wiring the ERFVII transcription factor RAP2.12 to PHD-mediated degradation in Arabidopsis. To this end, we uncoupled RAP2.12 from the PCO-branch of the Arg/N-degron pathway by substituting its N-cys-degron (aa 2-13) with the HIF_ODD_ peptide (**Fig. 3A-B**). This construct, controlled by the endogenous *RAP2.12* promoter, was expressed together with the *PHD3* and *SLY1-2-VHL* modules in a pentuple *erfVII* mutant^29^. We applied the same strategy used for O_2_ratio to generate independent lines expressing HIF_ODD_-ΔRAP2.12 and SLY1-2-VHL and super-transformed them with a *35S:PHD3* construct (**Supplementary Fig. 7A**). In this case, we used the promoter of the Arabidopsis gene *PCO4*^44^ to control *PHD3*, to ensure similar expression levels of the O_2_ sensing modules in the native plant mechanism and the synthetic mammal-inspired system. HIF_ODD_-ΔRAP2.12 stimulated the constitutive activation of the hypoxia marker genes *PCO1* and *SAD6;* co-expression of *PHD3* (particularly in Line #2) reversed this phenotype in normoxia, but was counteracted by hypoxia (**Supplementary Fig. 7B**). Altogether, this suggests that the engineered device is effective in controlling ERFVII activity. As a control, we separately introduced in the *erfVII* background a full RAP2.12 coding sequence, under the control of its own promoter. We tested the stability of the two RAP2.12 versions in a hypoxia time course by western blot, taking advantage of a fused double human influenza hemagglutinin (2xHA) tag. Wild type RAP2.12 was barely detectable under normoxic conditions and transiently stabilised after 1-2 h hypoxia (**Fig. 3C, Supplementary Fig. 8A**). The HIF_ODD_ version of RAP2.12 was more abundant than the native version under normoxia and further increased under hypoxia, but decreased after 4 h (**Fig. 3D, Supplementary Fig. 8B**). Remarkably, its stability also increased over time when plants were kept in the dark under aerobic conditions (**Fig. 3D, Supplementary Fig. 8B**).

**Figure 3.**
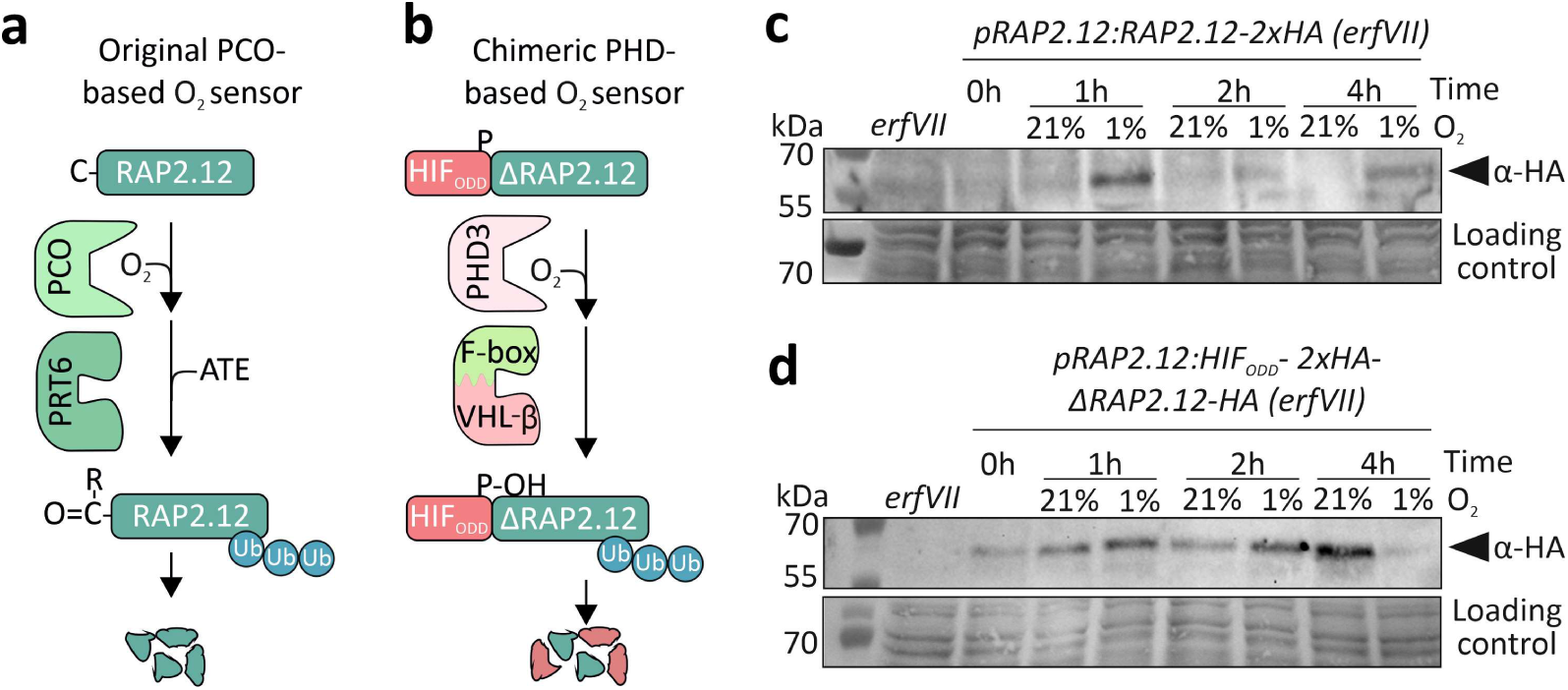
Complementation of the hypoxia-insensitive *erfVII* mutant by synthetic PHD-based oxygen sensing. **a,** A RAP2.12 fusion, used to complement the quintuple *erfVII* mutant, is controlled by the native PCO/N-degron pathway. **b,** Complementation of the *erfVII* mutant with a synthetic RAP2.12 version controlled by the synthetic O_2_ sensing device. Here, the N-terminal 13 aa were removed from RAP2.12 (ΔRAP2.12), to bypass PCO regulation, and substituted with HIF_ODD_. Under normoxia, HIF_ODD_-ΔRAP2.12 is expected to undergo PHD-dependent degradation. **c,** Immunodetection of RAP2.12 and **d,** of HIF_ODD_-ΔRAP2.12 following a treatment with 1% O_2_ for 0 to 4 hours. Ponceau staining of total proteins is shown as the loading control. The *erfVII* background (1% O_2_, 2 h) is shown as a negative control for both transgenic lines.

We decided to test whether the PHD-dependent version of RAP2.12 can regulate gene expression in response to hypoxia, similarly to what happens when this transcription factor is controlled via PCO and the N-degron pathway. We first explored this scenario by developing dynamic, ordinary differential equations (ODE)-based models to predict the expression dynamics of Hypoxia Responsive Genes (HRG) in either the PHD/VHL- or the N-degron pathway-based oxygen sensing systems. In these equations, we incorporated both Michaelis-Menten Kinetics and Mass Action laws to describe the gene expression and biochemical reaction network processes (**Supplementary Fig. S9A-B)**. The two models predicted early HRG activation upon exposure to 1% O_2_ in both the ‘wild type’ scenario, where RAP2.12 is under the control of the N-degron pathway, or the system where RAP2.12 is regulated by PHD and VHL- (**Fig. 4A**). However, the dynamic range of the latter was predicted to be much smaller, due to constitutively higher expression levels of the hypoxia reporter gene under aerobic conditions (time 0, **Fig. 4A**). We set out to test this prediction by comparing the induction of hypoxia marker genes in the two RAP2.12 complemented genotypes Arabidopsis lines described in **Fig. 3C** and **3D**. First, we measured the expression of the hypoxia marker gene(s) *PCO1* and *SAD6* in the *erfVII* mutant under normoxia (21% O_2_) and hypoxia (1% O_2_) in a 4 h time course. The mutant showed little residual capacity to elicit the expression of the anaerobic genes, as previously reported (**Fig. 4B** and **Supplementary Fig. 10A**)^45^. Both complementation strategies, instead, caused stronger expression of the reporter genes with induction dynamics similar to those predicted by our models (**Fig. 4B, Supplementary Fig. 10A**). We further compared the two hypoxia responsive devices in an experimental setup where plants were exposed to a range of subatmospheric O_2_ concentrations for 2 h. We observed that, while hypoxic gene expression was already induced at 10% O_2_ when under control of the N-degron pathway, the PHD system was only able to significantly increase RAP2.12 activity at 1% O_2_ (**Fig. 4C** and **Supplementary Fig. 10B**). Finally, we compared the tolerance to the hypoxic stress of plants equipped with the two alternative O_2_-sensing systems. We subjected 1-week old plants to 1% O_2_ for 7 days, under photoperiodic conditions. Both O_2_-sensing strategies were associated with decreased lethality as compared to the *erfVII* mutant, although the PCO-regulated RAP2.12 conferred better fitness than the PHD-regulated version (**Fig. 4D-E**). As previously observed (Shukla et al. 2019), higher RAP2.12 abundance in the latter genotype reduced primary root growth, both under aerobic and hypoxic conditions (**Fig. 4F**).

**Figure 4.**
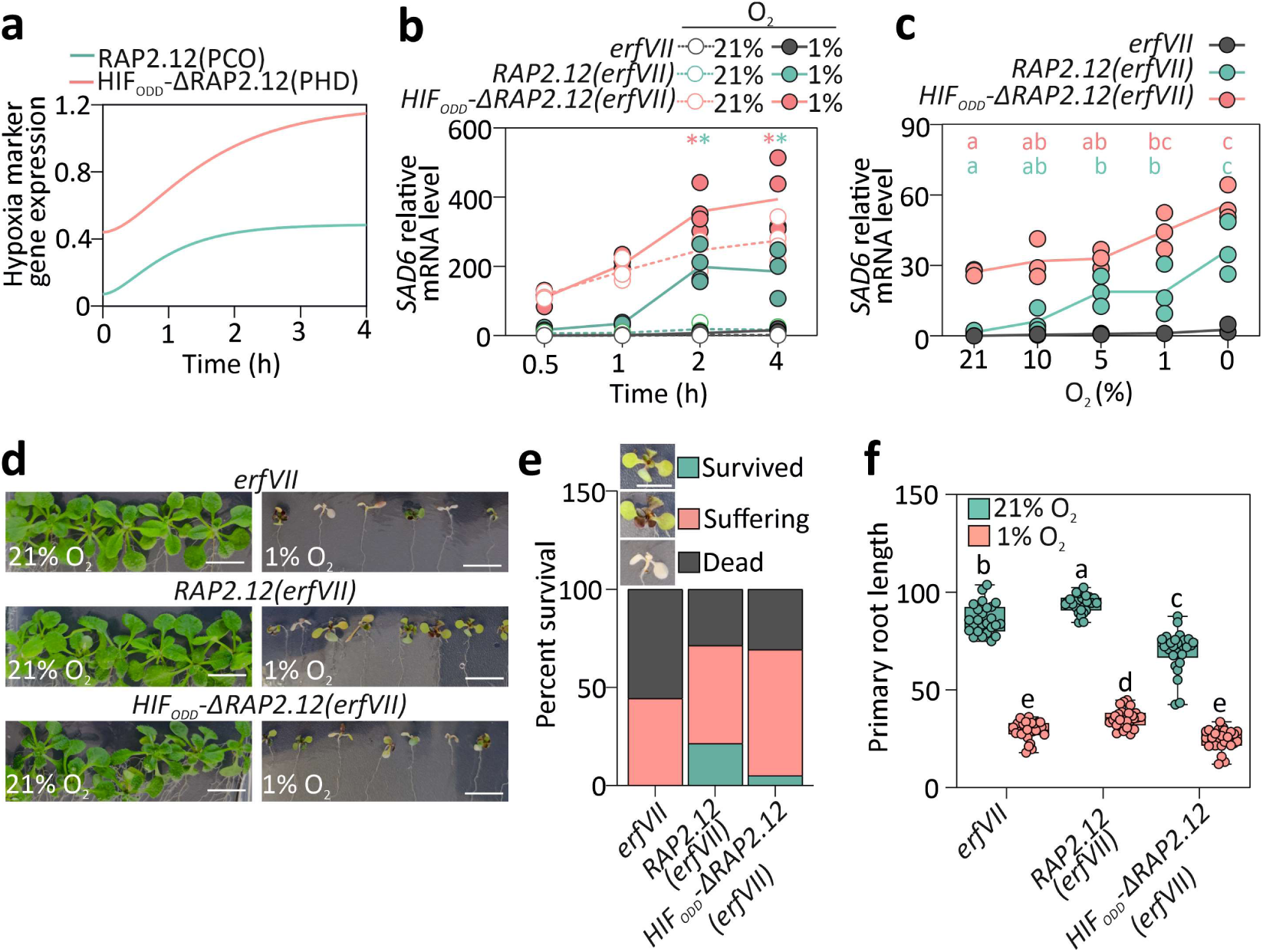
Functional characterization of native and synthetic O_2_ sensing mechanisms. **a,** Mathematical model prediction of the induction of a generic hypoxia responsive gene by RAP2.12 under the control of either the angiosperm O_2_ sensing mechanism or the mammalian-inspired synthetic sensing pathway. **b,** Expression of the hypoxia marker gene *SAD6* in 7-day-old Arabidopsis *erfVII* seedlings and complementation lines expressing RAP2.12-HA (teal) or HIF_ODD_-ΔRAP2.12-HA (pink) grown under normoxia and then subjected to 21% or 1% O₂ for 0.5 to 4 hours. Statistical significance of hypoxic gene expression relative to 21% O_2_ controls is denoted by asterisks. **c,** Dynamics of *SAD6* marker gene expression in response to different O₂ levels in *erfVII* plants expressing RAP2.12-HA under control of the PCO or PHD pathways in an *erfVII* background. Letters indicate statistical significance of hypoxic treatments compared with 21% O_2_ within each genotype, analysed independently by one-way ANOVA and followed by Tukey’s post-hoc test (p ≤ 0.05). **d,** Hypoxia tolerance assay of *erfVII* mutants and complemented lines expressing RAP2.12-HA or HIF_ODD_-ΔRAP2.12-HA. Seven-day-old seedlings were grown on agar plates and exposed to 1% O₂ for seven days, followed by a seven-day recovery period. Images were taken at the end of the recovery period. **e,** Quantification of plant fitness after hypoxic treatment characterized categorised as surviving, suffering or dead (n ≥36). Inset images illustrate representative status for categorisation and quantification. **f**, Primary root length measurements of seedlings from **d,** at the end of the seven-day hypoxia treatment. Letters indicate statistical significance, with two-way ANOVA and Tukey’s post-test (p ≤ 0.05, n≥27).

### Exploiting PHD-dependent proteolysis to engineer adaptive responses to flooding in plants

Having established that a metazoan-like mechanism for hypoxia sensing can successfully drive transcriptional responses in plant cells, we explored whether this can be connected to establish or enhance responses that support plant fitness under oxygen limitation. Reduced O_2_ availability commonly occurs because of soil waterlogging or whole plant submergence^25^. Flood adapted species have evolved two contrasting strategies: those frequently experiencing long-term and shallow submergence enhance stem and/or petiole elongation to escape the water level and thus maintain gas exchange, those instead exposed to deep or short-lasting floods rather suppress organ elongation and, in this way, reduce carbohydrate consumption, a strategy defined as ‘quiescence’^46^. We attempted to promote either response through the newly established PHD-dependent oxygen sensing system. The Arabidopsis Columbia-0 ecotype exhibits modest organ elongation underwater^47^. To enhance its escape syndrome, we decided to modulate the abundance of the transcription factor Phytochrome Interacting 4 (PIF4), the key regulator of petiole and hypocotyl elongation under shading and warm temperature^48,49^. We thus generated a hypoxia-inducible version of PIF4 made by its full coding sequence synthesized in frame with the HIF_ODD_ peptide. To minimise interference from other signalling pathways, we selectively mutated specific PIF4 residues whose phosphorylation in response to light, temperature and hormones is known to stimulate the proteasomal degradation of the protein ^50^ (**Fig. 5A**). Here, petiole elongation was expected to happen only under hypoxia, due to PIF-HIF proteolysis along normoxic plant growth (**Fig. 5B**). We sequentially transformed Col-0 plants with two different T-DNAs, one carrying both the *SLY1-2-VHL* and *pUBQ:PIF4-HIF_ODD_* expression cassettes, and a second bearing the *PHD3* cassette, according to the strategy adopted before (**Supplementary Fig. 5B and 7A**). *PIF4-HIF_ODD_* over-expression with a *UBQ10* promoter caused an elongated petiole phenotype that could not be inhibited by *PHD3*, even though driven by an equally strong promoter (**Supplementary Fig. 11**). We therefore resorted to use the ubiquitous, but moderately active, promoter of the *ROTUNDA3* (*RON3*, *At4g24500*) gene. We selected two independent lines for which we could segregate the *PDH3* transgene out and used these to measure hypocotyl (**Supplementary Fig. 12A-B**) and petiole length (**Supplementary Fig. 12C**). Expression of *pRON3:PIF4-HIF_ODD_* significantly stimulated growth of both organs, a phenotype that was reversed, at least in part, by co-expression of *PHD3* (**Supplementary Fig. 12C**). This indicated that our strategy to adopt PHD- and VHL-dependent degradation of developmental regulators is successful to control organ elongation. Finally, we tested whether environmental hypoxia could relieve the repression imposed by PHD3 and SLY1-2-VHL on the chimeric PIF4-HIF_ODD_. To do so, we submerged 15 days old plants grown in transparent cubic vessels with distilled water and measured petiole growth after 4 days. The wild type exhibited moderate elongation, which was insufficient to push the leaves near the water surface (**Fig. 5B-C**). Instead, petioles of the transgenic plants grew twice as much, bringing blades closer to the atmosphere (**Fig. 5B-C**). This result confirmed the possibility to adopt the mammal-inspired hypoxia sensing system to control growth in response to changes in ambient O_2_ availability.

**Figure 5.**
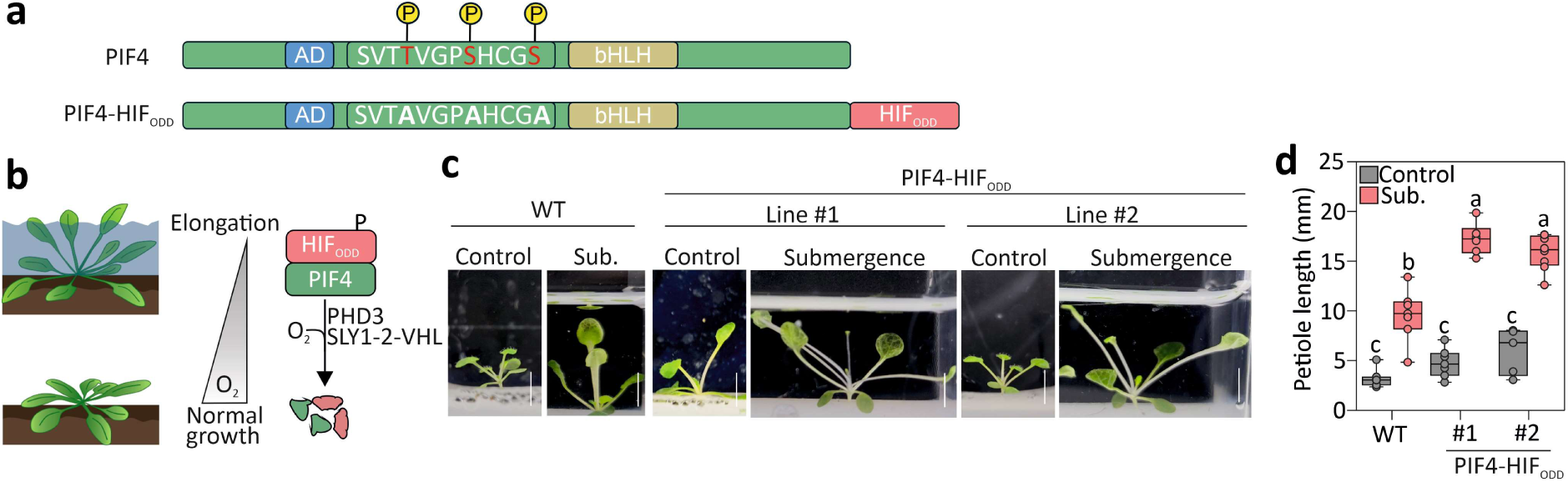
PHD and VHL-dependent control of petiole elongation under flooding induced hypoxia. **a,** Design of a PIF-HIF chimeric factor. Modifications applied to PIF4 to uncouple it from light, temperature and hormone regulation and rewire it to the mammalian-inspired synthetic O_2_ sensing system devised in this study. **b,** Predicted regulation of PIF4-HIF_ODD_. Under flooding-induced hypoxia, the PIF4-HIF protein is stabilized, promoting petiole elongation. Under normoxic conditions and with active PHD3, PIF4-HIF_ODD_ is targeted for degradation via SLY1-2-VHL. **c,** Phenotype of 2-week-old wild-type and PIF4-HIF_ODD_ plants treated for 4 days with darkness (control) or submergence (sub.). **d,** Quantification of petiole elongation from the plants in (**c**), showing that PIF4-HIF_ODD_ plants elongate their petioles in response to flooding. Letters indicate statistical significance, with two-way ANOVA and Tukey’s post-test (p ≤ 0.05).

## Discussion

With this work, we demonstrated the feasibility of engineering an orthogonal hypoxia responsive system for plant cells inspired by the O_2_-dependent degradation of the mammalian HIF proteins (**Fig. 1B**). We showed that this system is applicable to control stability, and thus activity, of different proteins, resulting in the possibility to modulate different outputs in an O_2_ sensing system. This approach aligns with those aimed to orthogonally control protein stability in mammalian and plant cells^51–53^. Successful engineering of selective proteolysis has been achieved with proteolysis- and autophagy-targeting chimera^54,55^, strategies that hold great potential for therapeutic applications and more recently explored also for agricultural applications^56^. Our exercise diverges from these in its aim to connect the output to a stimulus, cell O_2_ concentration, which is affected by both endogenous and external factors.

The process of design, test and optimization of the components required for this molecular device to perform the desired function highlighted old, and revealed new, opportunities and limitations of applying the synthetic biology framework to complex multicellular organisms. First, we confirmed that single cell systems, such as transiently transformed protoplasts, can be effectively used to test the performances of genetic circuits (**Fig. 1C-F and 2A-C**), before successful export to stable transformation in whole plants (**Fig. 3-5**). For example, our system performed similarly in both contexts, when it was used to control the stability of luciferase reporters (**Supplementary Fig. 2B and Fig. 2D**). Differences in output dynamic range seemed to be associated with the kind of protein used as reporter, as highlighted in our applications of the system to generate low oxygen reporters based on luminescence or fluorescence (**Fig. 2B,D** and **F**).

One of the main challenges faced in this exercise of biological engineering was the difficulty to estimate, and account for, the heterogeneous behavior of each component in a multicellular system. For example, cis-acting regulatory elements, such as promoters and regulatory mRNA untranslated regions, are strongly affected by their context: e.g. the chromatin state and endogenous cognate components (transcription factors and small RNAs) whose abundance can differ greatly between cell types. As one of the few attempts to implement multi-components genetic circuits in plants, it is interesting to observe the strong effect that alternative elements produced on the output (**Supplementary Fig. 5C**). This consideration becomes even more relevant when the genetic circuit relies on the control of protein stability, a process for which the characterization of components is more limited than transcription and translation. Such a limitation highlights the need of the community to produce quantitative characterisation of degradation determinants. Despite these constraints, the success of our strategy paves the way for further optimisation and implementation. Our attempts to alter output dynamics using structure-inspired PHD mutations that impact the catalytic site and neighboring residues did not succeed (**Supplementary Fig. 6D-E**), highlighting the difficulties in increasing the sensitivity of dioxygenases to hypoxia while maintaining high activity under aerobic conditions. The PHDs’ gas tunnel is a promising alternative target to test in the future, based on the enhanced hydroxylation efficiency achieved in mammalian cells^57^.

Transfer of PHD-based oxygen sensing from animal cells to plants interfaces with both fundamental and applied research. From a fundamental perspective, it allows experimental testing of the evolution of O_2_ sensing in eukaryotes and its diversification between plants and animals. Intriguingly, both kingdoms adopted proteostatic control of transcription factors, on top of which additional regulatory mechanisms insist, to reprogram cells to cope with reduced oxygen availability. While it has been speculated that this is a requirement and the optimal solution for the multicellular organization that involves stem cells^17^, it is puzzling that animals and plants diverged so much when their last common ancestor likely contained the biochemical functions to perform both N-cys-degron and VHL-dependent degradation. Successful control of transcriptional responses rewiring endogenous transcriptional regulators to PHD-primed degradation in Arabidopsis allows us to exclude intrinsic metabolic of signaling interference that would prevent this mechanism from working. This study also revealed the existence of additional regulation exerted on RAP2.12, which determines the transiency of its accumulation in hypoxia, and its increase in the dark (**Figure 3C-D**, **Supplementary Fig. 8A-B**). Similar, although slower, dynamics have been reported for the ERFVII RAP2.3^29^. Remarkably, target gene expression correlated with the protein abundance of native and synthetic RAP2.12 at early timepoints in the comparison between aerobic and hypoxic plants. This was not observed at later time points, which is likely explained by additional post-translational modifications, including phosphorylation that controls RAP2.12’s nuclear localization and transactivation capacity ^58,59^. N-degron dependent control of HIF stability, upstream of PHD regulation, has been proposed recently^60^. While it has been shown that N-cysteine oxidation determines substrate stability with similar O_2_-sensitivity to prolyl-hydroxylation in animal cells^61^, our swap exercise of O_2_ sensing system in plant cells hinted at higher proteolytic efficacy for the N-degron pathway when compared with VHL-promoted degradation. It is tempting to speculate that this might reflect the need to switch on adaptive responses at different O_2_ levels.

In summary, we successfully generated a PHD- and O_2_-dependent degradation of selected proteins through the engineering of a chimeric F-box protein. By adding the HIF_ODD_ fragment to different reporters and effector modules, we demonstrated the possibility to monitor cell oxic states as well as to drive advantageous developmental traits for submergence tolerance. From the applied perspective, wiring the synthetic O_2_ sensing system to control petiole elongation in Arabidopsis (**Fig. 5**) is a milestone in the engineering of adaptive responses to environmental cues. While it remains to be established whether promotion of fast petiole elongation can be useful in specific agroecosystems, it is possible to connect the novel O_2_ sensing mechanism to repressors of petiole elongation, thus favoring the quiescent syndrome similar to flash flood-tolerant rice varieties. The highly transgenic nature of the system prevents its immediate application in agriculture in many countries; nevertheless, it demonstrates feasibility and potential for synthetic biology to improve crops’ ability to cope with abiotic stresses, including those exacerbated by climate change. While we here conveniently used Arabidopsis as a testbed, given the compatibility of molecular components and the availability of simple transformation and selection protocols, the chimeric oxygen sensing system is likely exportable to many other crops, including monocots.

## Online Methods

### Plant materials and growth conditions

The *Arabidopsis thaliana* Columbia-0 (Col-0) ecotype was used as the wild-type background in all the experiments. The *erfVII* mutant^29^, which carries T-DNA insertions in the genes *At1g72360*, *At2g47520*, *At1g53910*, *At3g14230* and *At3g16770,* was provided by Michael J. Holdsworth. Plants were either grown in soil on a peat:perlite 3:1 mixture, or in sterility on agarized (8 g l^−1^) MS half-strength medium supplemented with 1% sucrose, after sterilization with 70% ethanol and 10% commercial bleach solution and 5 washes with sterile distilled water. In both cases, seeds were vernalized at 4°C for 48 h in the dark and then germinated at 22°C day/18°C night with a photoperiod of 12 h. The plant material was subjected to short-duration low-oxygen treatments as specified in the text and figure legends by flushing a mixture of pure nitrogen (N_2_) and compressed air to reach the desired O_2_ concentration inside hermetic boxes. For treatments longer than 24 h, we instead used a Hypoxic Workstations (Whitley). For submergence treatments, plants grown in sterility in magenta boxes were submerged with water at an approximate depth of 3 cm. The treatment began at the end of the day for a period of 72 hours.

### Generation of transgenic plants

*A. thaliana* plants were transformed with *Agrobacterium tumefaciens* applying the floral-dip method^62^. Selection of transformed plants was carried out using appropriate antibiotics or herbicides or fluorescent seed selection. Transgene presence was assessed by PCR.

### *In silico* protein complex assembly

The structure of the Cul1-Rbx1-Ask1-TIR1 complex was predicted based on atomic coordinates from the Protein Data Bank (PDB) entries 2P1Q and 2P1O (10.1038/nature05731) which include the TIR1 and ASK1 protein structures, and AlphaFold^63^ entries AF-Q94AH6-F1-v4 and AF-Q940X7-F1-v4, representing the Arabidopsis Cullin1 and Rbx1 structures, respectively.

Similarly, the Cul1-Rbx1-Ask1-SLY1 complex structure was predicted using the PDB entry 2P1Q containing the Ask1 protein structure, alongside AlphaFold entries AF-Q94AH6-F1-v4, AF-Q940X7-F1-v4 and AF-Q9STX3-F1-v4 which correspond to Cul1, Rbx1 and SLY1, respectively. AlphaFold models AF-Q94AH6-F1-v4, AF-Q940X7-F1-v4 and AF-Q9STX3-F1-v4 were retrieved from the AlphaFold protein structure database. Structural predictions of the Cul1-Rbx1-Ask1 complex with TIR1 and SLY1 were carried out using the full Cul1–Rbx1–Skp1–Skp2 complex as template (PDB: 1LDJ^64^).

The structure of the SCF complex Cul2-Rbx1-EloB-EloC-VHL-HIF_ODD_ was predicted using atomic coordinates 1LQB^65^ and 1VCB^63^, which include ELOB, ELOC, VHL and HIF_ODD_, and 5N4W^66^, which provides the structure for VHL, ELOB, ELOC, Cul2 and Rbx1. Structural alignment and figure rendering were performed using VMD and PyMol software.

### DNA cloning and assembly

Transcriptional units were either cloned from genomic DNA, cDNA or *de novo* synthesized by GeneArt (Thermo-Fisher Scientific). Protein-coding sequences were codon optimized using the EMBOSS Backtranseq online tools^67^. A full list of synthetic sequences and plasmids used in this work is provided in **Supplementary Table 1** and **2**, respectively. DNA fragments provided of a 5’-CACC additional sequence or flanked by restriction sites were initially sub-cloned either in the pENTR/D-TOPO^®^ vector, to be recombined into destination vectors using Gateway™ LR Clonase™ II Enzyme mix (catalog number 11791020, Thermo-Fisher Scientific), or in the pCR^™^2.1-TOPO^®^-TA vector (Thermo-Fisher Scientific), to be inserted into expression vectors via a restriction-ligation strategy, respectively. Plasmid maps were generated with Serial Cloner (Serial basic). A full list of primers used for cloning is provided in **Supplementary Table 3**.

For protoplast transformation, the 35S:FboxTIR1-VHL, 35S:HIF_ODD_-GAL4DBD-RAP2.12AD, 35S:SLY1-VHL, 35S:SLY1-2-VHL and 35S:FboxSLY1-10-VHL plasmids were generated from synthetic coding sequences, purchased as DNA strings from GeneArt (Thermo-Fisher Scientific), further ligated in the pENTR/D-TOPO^®^ vector (Thermo-Fisher Scientific) and recombined into the p2GW7^68^ vector via Gateway™ cloning. The 4xUAS:Fluc plasmid has been described in Bui et al. (2015)^37^. The 35S:PHD3 plasmid has been described in Iacopino et al. (2019)^23^

The O_2_ratio construct was produced by overlapping PCR. A 5’-terminal 3xHA-Renilla-UBQ fragment was amplified with the primers 3xHA-Fw and UBQ-Rv using the C-DLOR plasmid^69^ as template, while the fragment bearing the Firefly Luciferase CDS fused to the HIF_ODD_ was amplified with the primers UBQ_Luc-Fw and Luc_HIF_Rv using the 4XUAS:Fluc plasmid^37^ as a template. The Luc_HIF_Rv primer was designed to include the full HIF_ODD_ coding sequence. The two fragments were joined together with a second PCR using the oligonucleotides 3xHA-Fw and Luc_HIF_Rv. The resulting PCR product was cloned into the pENTR/D-TOPO^®^. The construct 35S:O_2_Ratio/pUBQ:SLY1,2-VHL used for Arabidopsis infiltration was generated in three steps. First, a synthetic construct consisting of the Arabidopsis UBQ10 promoter^70^, flanked by the SacI and AfeI restriction sites, and followed by the 35S CaMV terminator^71^, flanked by EcoRI and a further SacI restriction site, was ligated into the pK7WG2^68^ plasmid using the compatible SacI restriction site. Then,the SLY1-2 coding sequence was cloned in between the UBQ10 promoter and 35S CaM terminator using the AfeI and EcoRI restriction sites, to obtain a destination plasmid named pK7WG2/pUBQ10:SLY1-2. Finally, this destination was recombined with the entry plasmid containing O_2_ratio via Gateway™ cloning.

The NLS-O_2_ratio and O_2_ratio^2^ variants were generated by restriction/ligation cloning operated on the O_2_ratio entry vector. SacI and XbaI were used to remove a portion of the UBQ10 and FLuc sequence, and a DNA fragment designed to contain the excised regions and the additional HIF_ODD_ or NLS element was ligated using same restriction sites. The nanO_2_ratio^2^ entry vector was synthesized by GeneArt (Thermo Fisher Scientific). The nanO^2^ratio variants were generated by digesting the nanO_2_ratio^2^ entry vector using the SacI and XhoI restriction sites and ligating an identical DNA sequence devoid of the first HIF-COOD. To replace the CaMV 35S Omega leader with the 5’UTR region of the Arabidopsis *ADH1* gene, pK7WG2/pUBQ10:SLY1-2 was cut with StuI and SpeI and ligated with a synthetic DNA string containing the excised region of the 35S promoter followed by *ADH1* 5’-UTR, flanked by compatible restriction sites thereby turning the original vector into the new pK7WG35SUTR^ADH^/pUBQ10: SLY1-2 Gateway destination vector.

All O_2_ratio variants entry vector were then recombined using Gateway™ cloning in these two destination vectors to generate the binary expression vectors reported in **Supplementary Figure 5**.

The NLS-VENUS-HIF_ODD_ construct was purchased as DNA string from Geneart (Thermo-Fisher Scientific) and ligated into the pENTR/D-TOPO^®^. The p16:NLS-VENUS-HIF_ODD_/pUBQ: SLY1-2-VHL plasmid was generated by recombination of the NLS-VENUS-HIF_ODD_ entry with pK7WG2/pUBQ10:SLY1-2 and subsequent replacement of the 35S promoter. Specifically, the expression plasmid was cut with StuI and SacI restriction ezymes and ligated with a compatible p16 promoter sequence^43^ amplified using primers p16_Fw and p16_Rv and from a p16-FLIPnls43 template vector^72^.

The construct pRAP2.12:HIF_ODD_-2xHA-D13RAP2.12/pUBQ:SLY1-2-VHL was generated as follows: a synthetic DNA string containing the chimeric HIF_ODD_-2xHA-D13RAP2.12 coding sequence was ligated in pENTR/D-TOPO^®^ and recombined with the pK7WGpUBQ10:SLY1-2 destination vector. A 2kb-long upstream region of the *RAP2.12* promoter was amplified using NcoI_pRAP_Fw and NcoI_pRAP_Rv from Arabidopsis genomic DNA. The 35S promoter was then removed from the expression plasmid using NcoI restriction enzyme and substituted with the *RAP2.12* promoter fragment via ligation. Finally, the kanamycin plant selection marker was removed using ApaI and CpoI restriction enzymes, to be replaced with a fluorescent selection cassette. The pOLE_Fw and mCHERRY_Rv primers were used with a pKIR1.0^73^ template plasmid, to amplify an expression cassette consisting of the Arabidopsis oleosin promoter (pOLE1) and a downstream fusion sequence between the red fluorescent protein mKATE2 and the oleosin gene (*OLE1*). This fluorescent cassette, flanked by compatible ApaI and CpoI sites, was subsequently ligated with the backbone obtained before. The construct pRAP2.12:RAP2.12-2xHA was synthesized in an entry plasmid by GeneArt (Thermo-Fisher Scientific) and subsequently recombined into the pH7WG^68^ binary vector via Gateway™ cloning.

To generate the pPCO4:PHD3 expression vector, the PHD3 entry plasmid was recombined Gateway™ cloning in the plasmid pH7WG-promPCO4^44^. The expression plasmid was digested with ApaI and CpoI and an expression cassette consisting of the Arabidopsis At2S3 promoter driving GFP expression amplified from pFP91^74^ with primers At2S3_Fw and t35S_Rv and subsequently ligated. PHD3 mutations were obtained by site directed mutagenesis with PHD3Asp137His Fw and Rv, PHD3Asp137Glu Fw and Rv and PHD3Arg205Lys Fw and Rv. The PIF4-HIF_ODD_ were purchased as DNA string and ligated into the pENTR/D-TOPO^®^. The pSIC:PIF4-HIF_ODD_/pUBQ:SLY1-2-VHL plasmid was obtaining recombining the PIF4-HIS entry into the pK7WG2/pUBQ:SLY1-2 and replacing the 35S CaMV promoter using StuI and SpeI restriction site. The RON3 promoter was amplified by PCR using primers pAt4g24500_Fw and pAt4g24500_Rv using Col-0 genomic DNA as template and then ligated using compatible restriction sites.

### Protoplasts transformation and transactivation assay

Protoplasts isolation and transformation were performed as reported by Yoo et al.,^75^ with the modifications described in Iacopino et al.^23^. Firefly and Renilla luciferase activity was measured using the Dual-Luciferase® Reporter Assay System (E1910, Promega), Nanoluciferase and Firefly luciferase activity was measured using Nano-Glo® Dual-Luciferase® Reporter Assay System (N1610, Promega).

### Phenotypic analyses

Petiole and hypocotyl lengths were measured from images using ImageJ software^76^. Petioles were pressed between two layers of Scotch tape and scanned with an EPSON Perfection V750 PRO scanner at a 400 DPI resolution. Hypocotyl images were captured using a Nikon Coolpix P520 digital camera. Root lengths were measured from images scanned from plastic square plates using ImageJ software.

### Confocal imaging

Confocal investigations were performed using a Zeiss LSM800 confocal microscope. For sub-cellular localization studies, GFP fluorescence was excited with 488 nm laser light and collected with a 497-554 nm long-pass emission filter. Chlorophyll autofluorescence was excited at 633 nm and collected at 650–750 nm. Nuclei were stained with 1µg µl^−1^ 4’,6-diamidino-2-phenylidone (DAPI, Sigma-Aldrich), and fluorescence excited at 405 nm and collected at 410-470 nm. For Venus visualization in roots, plants were stained with 10 µg µl^−1^ propidium iodide (PI) (Sigma-Aldrich) cell wall stain. The roots were observed with 20x objective lens under ZEISS LSM 800 Laser Confocal Scanning Microscope. HIF-NLS-Venus seedlings were fixed with 4% paraformaldehyde and stained with ClearSee^77^ supplemented with 1 µl/ml SCRI Renaissance 2200 (SR2200). Venus fluorescence was excited with 488 nm laser light and collected with a 520-560 nm. PI was excited with 488 nm laser light and collected at 650-700 nm. Images were analyzed with the ZEN 2010 software (Zeiss).

### SDS-PAGE and immunoblotting

Total proteins were isolated from 1-week old seedlings grown on vertical plates using a buffer containing 50mM Tris (pH7.5), 0.1% w/v SDS and Protease Inhibitor Cocktail (cOmplete™, Mini, EDTA-free Protease Inhibitor Cocktail, Sigma-Aldrich; 11836170001). From leaf mesophyll protoplasts, total proteins were instead extracted using a buffer containing Laemmli and 5% v/v β-mercaptoethanol. Equal total protein amount (70 μg) were resolved by SDS-PAGE and then transferred to PVPF membrane using MiniTrans-Blot electrophoretic transfer cell (Bio-Rad). To detect the HA tag, membranes were probed with anti-HA primary antibody (Sigma-Aldrich; H3663) at 1:2000 dilution. To detect the GAL4 activation domain, membranes were probed with the monoclonal anti-GAL4 antibody [15-6E10A7] (ab135397, Abcam) at 1:1000 dilution. In both cases, membranes were then probed with HRP-conjugated anti-mouse secondary antibody (Sigma-Aldrich; 12-349) at 1:10000 dilution. Immunoblots were developed with SuperSignal™ West Pico PLUS Chemiluminescent Substrate (Thermo Fisher Scientific) using the iBright CL1500 Imaging System (Thermo Fisher Scientific).

### Plant RNA isolation and qPCR analyses

Total RNA was extracted from 100 mg of frozen-ground seedlings as described previously^32^. One microgram of total RNA was treated with DNase (Thermo Scientific™) to remove genomic DNA before retro-transcription using qPCRBIO cDNA Synthesis Kit (PCR Biosystems). Real-time quantitative PCRs were performed in 10 μl volume using a 2X Power SYBR™ Master Mix (Thermo Fisher Scientific™), 10 ng cDNA and 0.2 μM of specific reverse and complement primers for each gene to be tested. Thermal cycling and fluorescence acquisition was carried out with StepOnePlus™ Real-Time PCR System. Ubiquitn10 (AT4G0532) was used as housekeeping genes for Arabidopsis analysis. A full list of the primers used for qPCR is included in **Supplementary Table 3.** Relative expression of each individual gene was calculated using the 2^−ΔΔCt^ method^78^.

### Statistical analysis

Ordinary two-way and one-way analysis of variance (ANOVA) and multiple comparisons for statistical differences were performed with GraphPad Prism 9 for Windows 10.

### Mathematical modelling of RAP2.12/PCO4 and HIF-RAP2.12/PHD kinetics

#### Model description and assumption

The two models that were developed to describe the expression dynamics of Hypoxia Responsive Genes (HRG) in either the PHD/VHL- or the N-degron pathway-based oxygen sensing system consist of two coupled Ordinary Differential Equations (ODEs). Within these equations, we incorporated both Michaelis-Menten Kinetics and Mass Action laws to describe the gene expression and biochemical reaction network processes (**Supplementary Tables 4 and 5**). The overall processes involving RAP2.12 expression, degradation, and downstream HRGs activation illustrated in **Supplementary Fig. S9**.

In our models, the RAP2.12 protein is synthesized at a rate k_1_ and degraded at a rate k_2_. In the *promRAP2.12:RAP2.12-2xHA* scenario, where RAP2.12 is the only ERFVII expressed in plant cells, the N-Cys-degron pathway promotes O_2_-dependent RAP2.12 degradation following Michaelis-Menten kinetics, with catalytic constant k_3_, and Michaelis constants Km_1_ and Km_2_. In this system, HRGs expression, which in our model is assumed to be exclusively driven by RAP2.12, follows Michaelis-Menten kinetics at a rate represented by k_4_ and Km_3_ and decay occurring at a rate of k_5_. In the HIF_ODD_-RAP2.12 system, we applied the same parameters for RAP2.12 protein synthesis and degradation as well as for HRG activation. However, in this chimeric system, HIF_ODD_-RAP2.12 undergoes O_2_-dependent degradation through enzymatic prolyl-hydroxylation, characterized by rates k_6_, Km_4_, and Km_5_. Since the production and degradation rates of RAP2.12 are unknown, we used the HIFα protein production and degradation rates from the literature. Similarly, the PHD concentration was set to 0.1 µM based on published models^79^. These assumptions do not affect the validity of our conclusions.

To simplify our model, we assume that all reactions occur in a homogeneous environment, without accounting for compartmentalization or specific microenvironments within the cell. We further assume that each reaction species—such as RAP2.12, PCO, PHD, O_2_, and mRNA—is evenly distributed throughout the system. To focus on the oxidation and hydroxylation processes, we simplify the model by assuming that RAP2.12 and HIF_ODD_-RAP2.12 are degraded instantly following N-Cys and Pro oxidation, respectively.

Additionally, for comparability, we assume identical concentrations of PCO and PHD and at steady state under normoxic conditions that characterise the initial timepoint in the simulation. These enzyme concentrations are assumed to remain constant after exposure to 1% hypoxia. In a short timeframe, this is not unrealistic since protein synthesis is strongly reduced under hypoxia^80^. By maintaining the same levels of PCO and PHD, we can compare the dynamic hypoxic responses based solely on their catalytic activities.

#### Non-linear fitting and simulation

We used MATLAB’s ‘Welsh’ and ‘Fair’ weight functions within the nlinfit function. The rate constants k_3_, Km_1_, and Km_2_ were re-fitted for the PCO-mediated degradation of RAP2.12, while k_6_, Km_4_, and Km_5_ were re-fitted for the PHD-mediated degradation of HIF_ODD_-RAP2.12. This three-dimensional nonlinear fitting, which is depicted in **Supplementary Fig. S9B**, was performed based on published kinetic data^21,81^. All parameters are listed in Table S4.

To simulate hypoxic responses, the initial concentrations of RAP2.12/HIF_ODD_-RAP2.12 and mRNA as inputs were obtained from steady-state simulations conducted under normoxic conditions (21% O_2_) for six hours (**Supplementary Table 6**). Subsequently, dynamic simulations of hypoxic responses were conducted under 1% O_2_ conditions for 200 minutes.

