## Supplemental Information for "Engineering prolyl hydroxylase-dependent proteolysis enables the orthogonal control of hypoxia responses in plants"

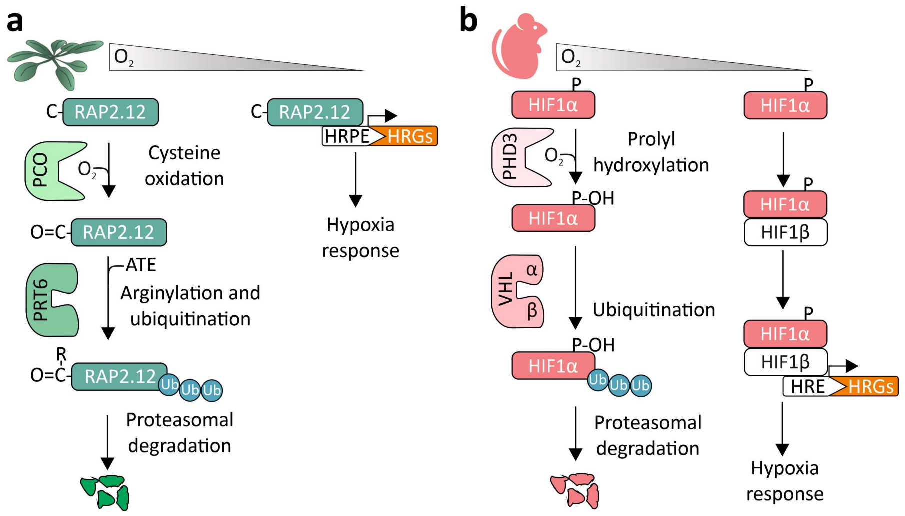


**Figure S1**: **$O_2$ sensing in vascular plants and in mammals. a**, Plant $O_2$ sensing based on the Plant Cysteine Oxidase (PCO)/N-degron pathway. After the removal of the initial Met by Met Aminopeptidases (MAP), the N-terminal Cys residue (C) of ERFVIIs is labelled for proteasomal degradation by the sequential action of PCOs, Arginyl-tRNA transferase (ATE) and the Proteolysis6 (PRT6) E3 ubiquitin ligase. Under hypoxic conditions, ERFVIIs are stabilised and induce the expression of hypoxia-responsive genes. **b**, Animal $O_2$ sensing based on the VHL pathway. In the presence of $O_2$, the α-subunit of the heterodimeric HIF-1 transcription factor is hydroxylated at two proline residues (P) by prolyl hydroxylases (PHDs). This modification marks HIF-1α for proteasomal degradation via the E3 ubiquitin ligase VHL. Under hypoxia, the stabilised HIF-1α forms a dimer with HIF-1β and activates the expression of target genes.

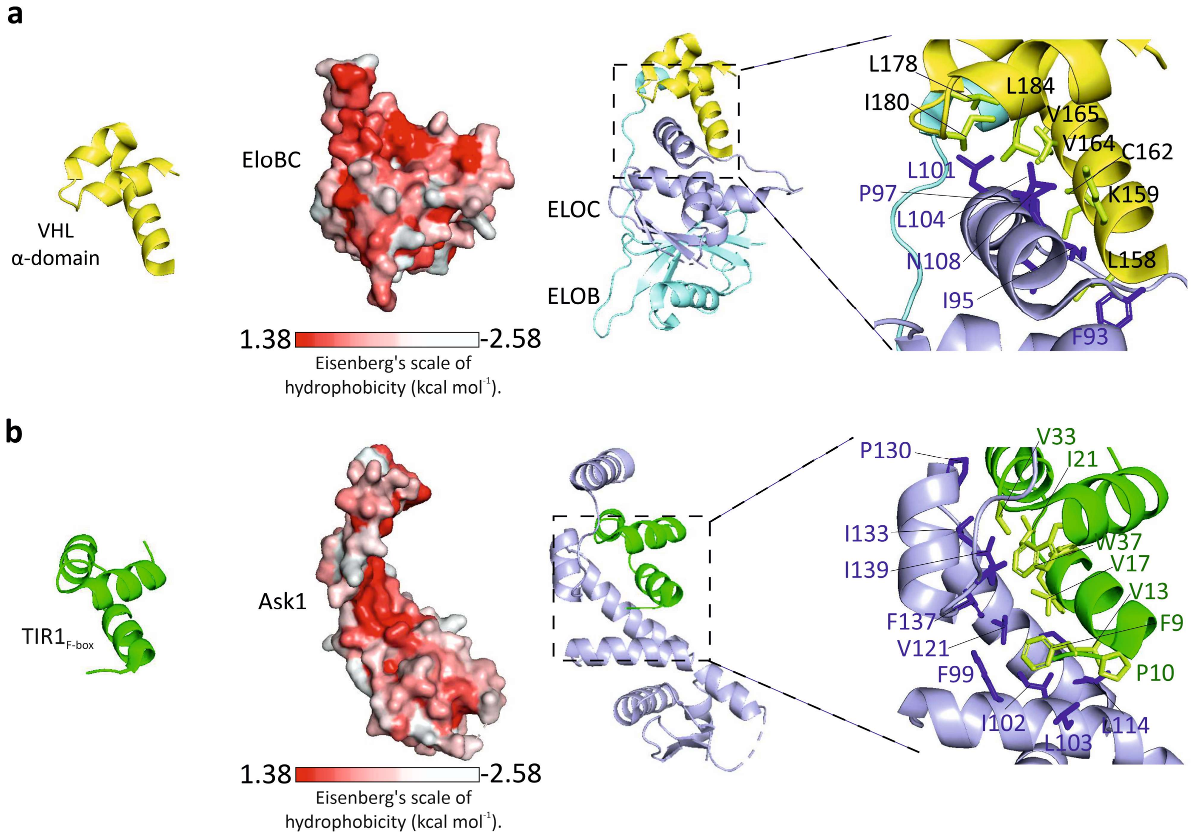


**Figure S2. Structural comparison of plant F-box domains and the α-domain of human VHL.** **a**, From left to right: cartoon representation of the VHL α-domain, surface representation of the Elongin B (ELOB) and ELOC heterodimer and cartoon representation of the complex containing the VHL α-domain and ELOB and ELOC. The inset provides a close-up view of the residues involved in the interaction. The VHL α-domain is coloured in yellow, ELOB and ELOC surface was coloured according to the Eisenberg's scale of hydrophobicity using the color_h script in PyMOL. [1]In this scale, red colour indicates hydrophobicity, from least hydrophobic (white) to most hydrophobic (red). In the cartoon representation on the right, ELOB and ELOC are coloured in blue and cyan, respectively. **b**, From left to right: cartoon representation of the F-box domain of Transport Inhibitor Resistant 1 (TIR1), surface representation of the Ask1 protein and cartoon representation of Ask1-$TIR1_{F\text{-}box}$ interaction with a close-up view of the residues involved. The $TIR1_{F\text{-}box}$ is coloured in green, Ask1 surface is coloured according to Eisenberg's scale of hydrophobicity[1] and the Ask1 cartoon is coloured in blue. TIR1 and ASK1 structures were retrieved from PDB: 2P1O[1]. The structures of VHL, ELOB and ELOC were obtained from PDBs 1VCB[2] and 1LQB[3].

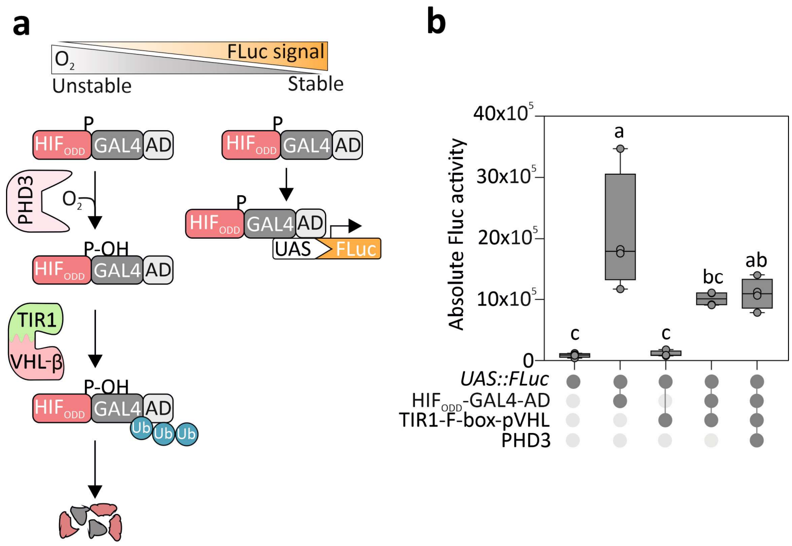


**Figure S3. Test of a TIR1-containing hypoxia-inducible $O_2$ sensor. a**, Schematic representation of the synthetic device. The human $HIF_{ODD}$ was equipped with the Gal4 DNA binding domain (DBD) end the Arabidopsis RAP2.12 activation domain (AD), while the VHL β-domains (VHL) was fused in frame with the F-box domain of TIR1 . In the presence of $O_2$, the O2-dependent degradation domain of $HIF_{ODD}$-GAL4-AD is hydroxylated by PHD3 and therefore recognised by VHL, which mediates its ubiquitination and subsequent proteasomal targeting. At low $O_2$ levels, instead, $HIF_{ODD}$-GAL4-AD can bind repeats of the GAL4 upstream activating sequence (4xUAS) in a synthetic promoter and induce the transcription of a firefly luciferase reporter (FLuc). **b**, Sensor output observed in Arabidopsis mesophyll protoplasts after transfection with different combinations of the effector modules $HIF_{ODD}$-GAL4-AD, $TIR1_{F\text{-}box}$ -VHL and PHD3 (empty circles, effector absent; dark circles, effector present). Letters indicate statistical differences ($p \leq 0.05$) calculated by a two-way ANOVA comparison followed by Tukey's post-test (n=4).

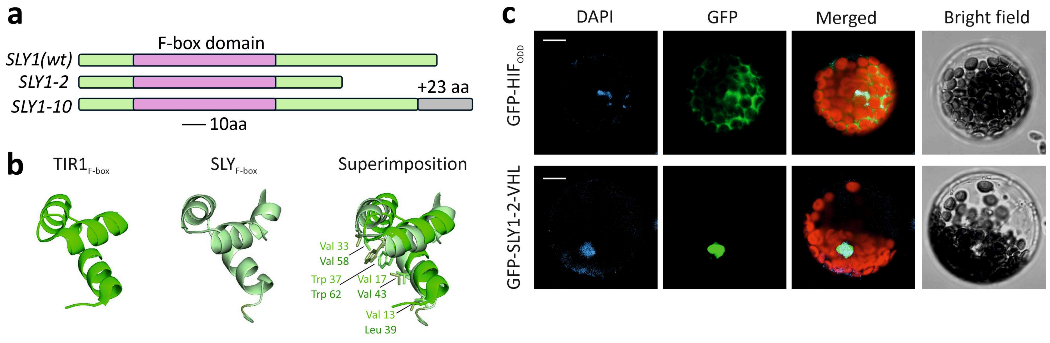


**Figure S4. Design of a SLY1-containing hypoxia-inducible $O_2$ sensor and localization of its effector modules. a,** Schematic representation of wild-type SLY1 and the two mutant variants SLY1-2 and SLY1-10. Both variants consist of SLY1 truncations, with an additional 23 aa-long domain at SLY1-10 C-terminal end. **b,** Comparison of the structure of the F-box domain of TIR1 and SLY1 and their superimposition showing the conservation of residues depicted in Supplementary figure 2B. **c,** Subcellular localization of the GFP-SLY1-2-VHL and GFP-$HIF_{ODD}$ constructs imaged in isolated Arabidopsis mesophyll protoplasts. DAPI staining marks the nuclei. In the merged image, the red colour is associated with chloroplast auto-fluorescence. Scale bar = 10μm.

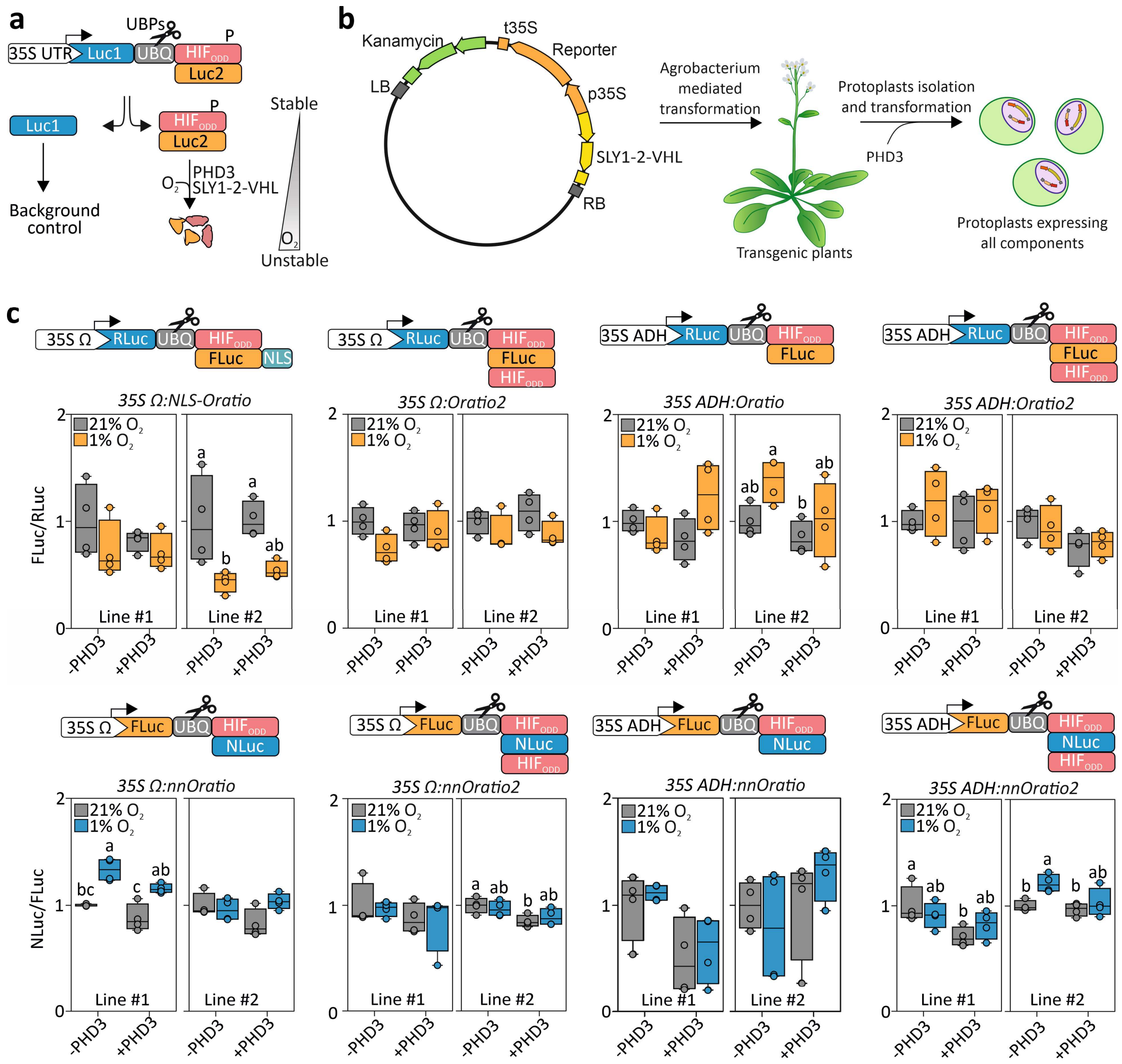


**Figure S5. Development and performance of a ratio-metric $O_2$ sensors in plants.** **a,** A polyprotein consisting of luciferase 1 (Luc1), UBQ10 (UBQ) monomer, and $HIF_{ODD}$ fused to a different luciferase (Luc2) is expressed in plant cells. Endogenous ubiquitin proteases (UBPs) co-translationally cleave the polypeptide at the c-terminus of UBQ10. The resulting LUC1-UBQ serves as a normalization reference while $HIF_{ODD}$-LUC2 is used to monitor $O_2$ levels. **b,** Strategy used to test variants of the synthetic $O_2$ reporter in protoplasts. Two stable transgenic lines were generated for each reporter and SLY1-2-VHL construct. Protoplasts were obtained from each line to test the effect of PHD3 expression on reporter activity under aerobic (21% $O_2$) and hypoxic (1% $O_2$) conditions. Such treatments were applied 12 h after protoplast transection and lasted 12 h. **c,** Relative output of different reporter variants tested using the strategy described in (B). FLUC2/FLUC1 ratio in the absence of PHD3 was set to 1 FLuc, firefly luciferase; RLuc, renilla luciferase; NLuc, nanoluciferase. 35S Ω, CamV 35S promoter with omega leader; 35S ADH, CamV 35S promoter with 5'-UTR sequence from the Arabidopsis *ADH1* gene; NLS, nuclear localization signal from *S. cerevisiae* Gal4. Letters indicate statistical differences ($p \leq 0.05$) determined by two-way ANOVA followed by Tukey's post-hoc test (n = 4).

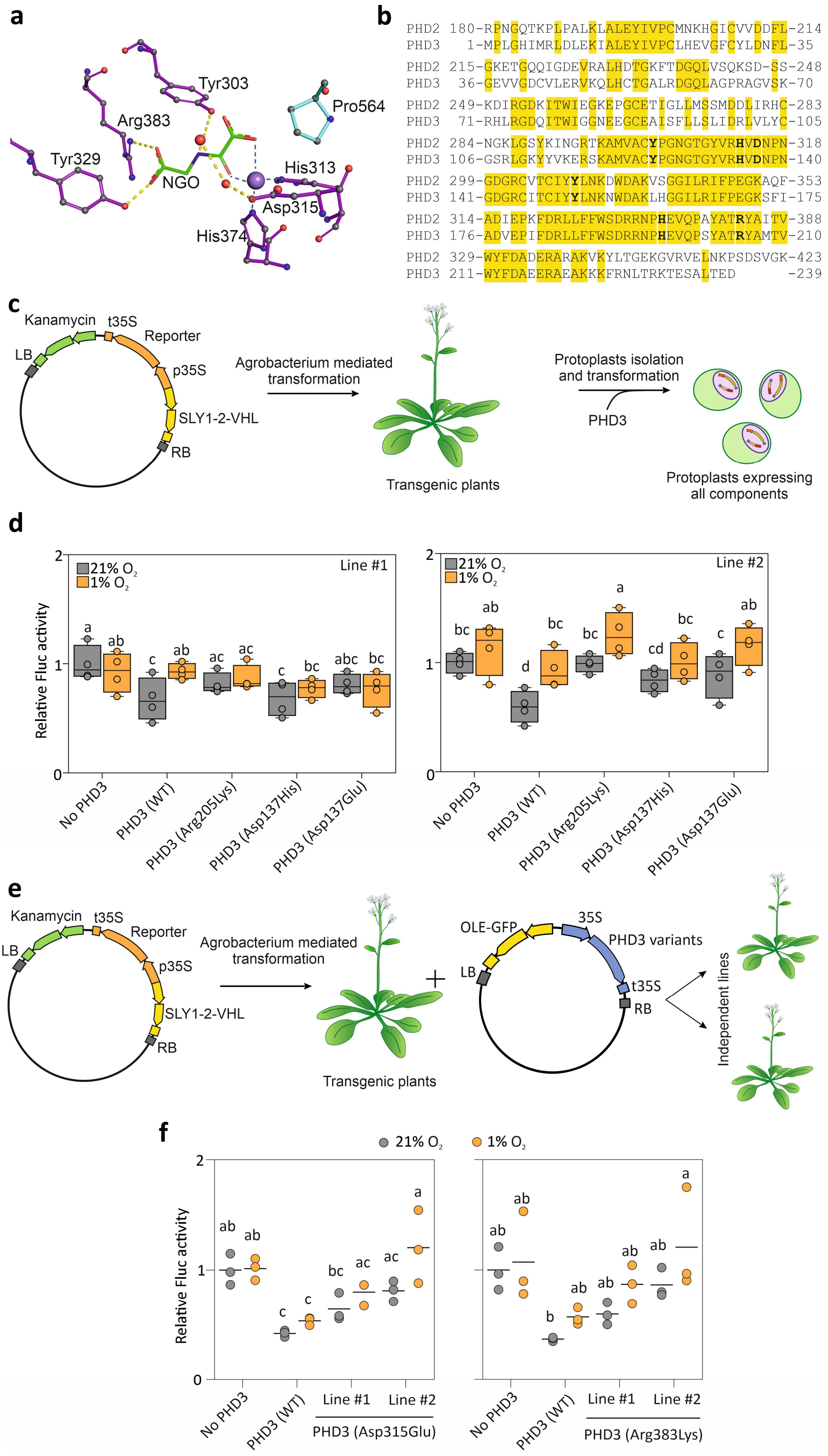
a
Tyr303
Pro564
Arg383
Tyr329
NGO
His313
Asp315
His374
b
PHD2 180-RPNGQTKPLPALKLALEYIVPCMNKHGICVVDDFL-214
PHD3 1-MPLGHIMRLDLEKIALEYIVPCLHEVGFCYLDNFL-35
PHD2 215-GKETGQQIGDEVRALHDTGKFTDGQLVSQKSD-SS-248
PHD3 36-GEVVGDCVLERVKQLHCTGALRDGQLAGPRAGVSK-70
PHD2 249-KDIRGDKITWIEGKEPGCETIGLLMSSMDDLIRHC-283
PHD3 71-RHLRGDQITWIGGNEEGCEAISFLLSLIDRLVLYC-105
PHD2 284-NGKLGSYKINGRTKAMVACYPGNGTGYVRHVDNPN-318
PHD3 106-GSRLGKYYVKERSKAMVACYPGNGTGYVRHVDNPN-140
PHD2 299-GDGRCVTCIYYLNKDWDAKVSGGILRIFPEGKAQF-353
PHD3 141-GDGRCITCIYYLNKNWDAKLHGGILRIFPEGKSFI-175
PHD2 314-ADIEPKFDRLLFFWSDRRNPHEVQPAYATRYAITV-388
PHD3 176-ADVEPIFDRLLFFWSDRRNPHEVQPSYATRYAMTV-210
PHD2 329-WYFDADERARAKVKYLTGEKGVRVELNKPSDSVGK-423
PHD3 211-WYFDAEERAEAKKKFRNLTRKTESALTED -239
c
Kanamycin
t35S
Reporter
LB
p35S
SLY1-2-VHL
RB
Agrobacterium mediated transformation
Transgenic plants
Protoplasts isolation and transformation
PHD3
Protoplasts expressing all components
d
Relative Fluc activity
21% O2
1% O2
Line #1
Line #2
No PHD3
PHD3 (WT)
PHD3 (Arg205Lys)
PHD3 (Asp137His)
PHD3 (Asp137Glu)
e
Kanamycin
t35S
Reporter
LB
p35S
SLY1-2-VHL
RB
Agrobacterium mediated transformation
Transgenic plants
OLE-GFP
35S
PHD3 variants
LB
t35S
RB
Independent lines
f
21% O2
1% O2
Relative Fluc activity
No PHD3
PHD3 (WT)
Line #1
Line #2
PHD3 (Asp315Glu)
PHD3 (Arg383Lys)

**Figure S6. Generation and testing of PHD3 variants. a,** Catalytic site of PHD2 human protein extracted from the PDB file 2G19[4] and readapted from (Tarhonskaya et al., 2014). **b,** Amino acid sequence alignment of PHD2 and PHD3. The conserved residues are highlighted in yellow. Bold font indicates residues represented in A. **c,** Strategy used to test expression of PHD3 variants in protoplasts stably expressing $O_2$ratio and SLY1-2-VHL. **d,** Sensor output in mesophyll protoplasts isolated from $O_2$ratio and SLY1-2-VHL expressing Arabidopsis plants and transformed with PHD3 expression plasmids. 12 h after transformation, protoplasts were either subjected to 6h of hypoxia (1% $O_2$) or normoxia (21% $O_2$). The FLuc and RLuc ratio in the absence of PHD3 was set to 1. Letters indicate statistical differences ($p \leq 0.05$) calculated by a two-way ANOVA comparison followed by Tukey's post-test (n=4). **e,** Strategy used to test efficiency of PHD3 variants in plants stably expressing $O_2$ratio, SLY1-2-VHL and different PHD3 variants. **f,** Sensor output in stable transgenic plants expressing $O_2$ratio module and SLY1-2-VHL super-transformed with binary plasmid containing PHD3 variants. One-week-old seedlings were either subjected to 6 h of hypoxia (1% $O_2$) or normoxia (21% $O_2$). The FLuc/RLuc ratio in the absence of PHD3 was set to 1. Letters indicate statistical differences ($p \leq 0.05$) calculated by a two-way ANOVA comparison followed by Tukey's post-test (n=3).

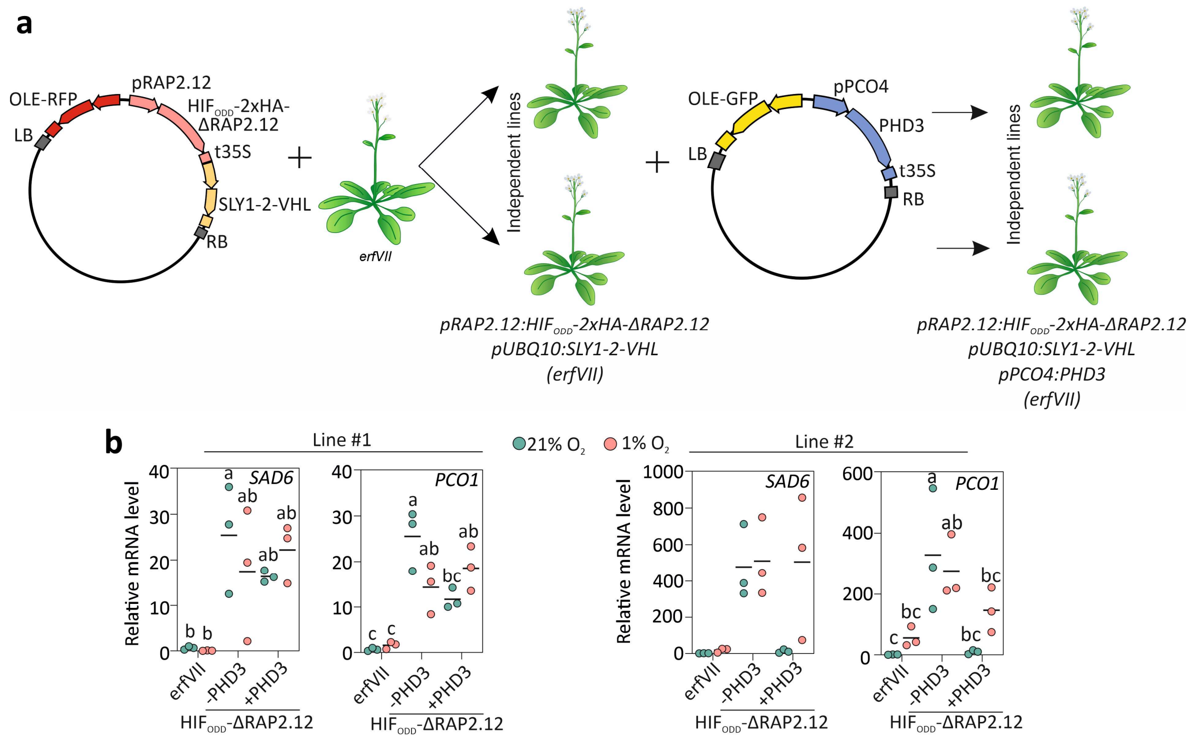


**Figure S7. Complementation of the *erfVII* mutant with the chimeric $O_2$ sensing system. a,** Two independent transgenic lines expressing HIF$_{ODD}$-ΔRAP2.12 and SLY1-2-VHL in the *erfVII* background were super-transformed with a binary vector containing *pPCO4:PHD3* to complete the $O_2$ sensing system. **b,** Two independent lines expressing HIF$_{ODD}$-ΔRAP2.12, SLY1-2-VHL and PHD3 were tested for PHD- and $O_2$-dependent expression of hypoxia marker genes *SAD6* and *PCO1*. Seven-day-old Arabidopsis seedlings were exposed to 21% or 1% $O_2$ for 2 hours. Letters indicate statistical significance evaluated by a two-way ANOVA followed by the Tukey's post-hoc test for multiple comparisons.

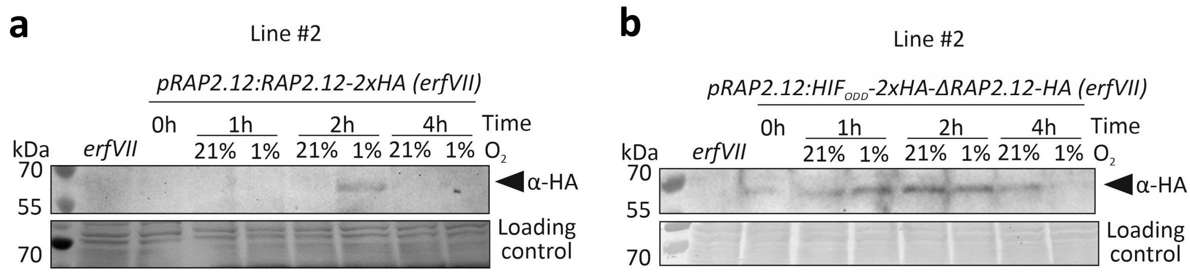


**Figure S8. Immunodetection of RAP2.12 in the *erfVII* background complemented with the original and chimeric oxygen sensing systems. a,** Immunodetection of RAP2.12-2xHA (line #2) under the regulation of the PCO degradation pathway following a treatment with 1% $O_2$ for 0 to 4 hours. The untransformed *erfVII* genotype (1% $O_2$, 2h) is included as a negative control. **b,** Immunodetection of $HIF_{ODD}$-ΔRAP2.12-2xHA (line #2) under aerobic conditions or after treatment with 1% $O_2$ for 0 to 4 hours, with the *erfVII* genotype (1% $O_2$, 2h) included as a negative control. Ponceau staining of total loaded proteins serves as the loading control for Western blots.

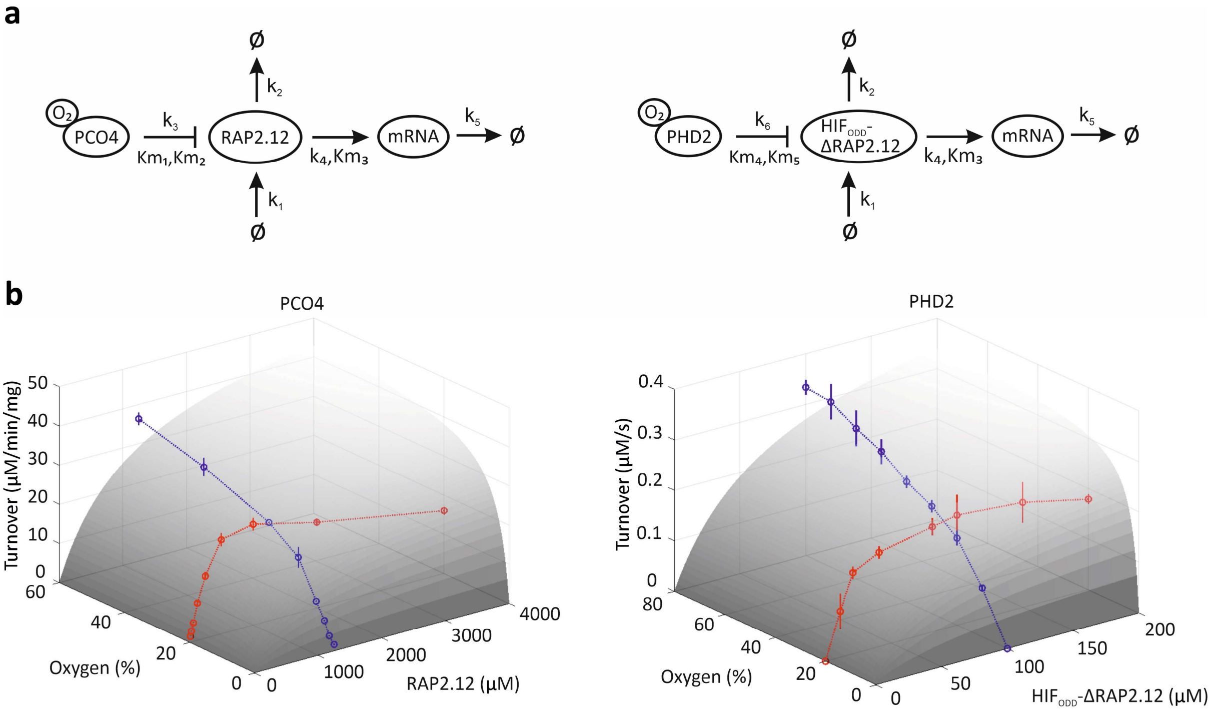


**Figure S9. Mathematical modelling of RAP2.12 abundance and target gene expression depending on $O_2$ levels. a,** Schematic representation of models for the endogenous (left) and the chimeric (right) $O_2$ sensing system tested in plant cells in this study. All model species including RAP2.12, HIF-RAP2.12, PCO4, PHD, $O_2$ and HRG mRNA are circled, and 'Ø' symbol stands for 'null'. **b,** 3D data visualization and nonlinear fitting for RAP2.12/PCO4 and HIF-RAP2.12/PHD kinetics. The 3D surface plot (grey) shows predicted turnover as a function of oxygen and substrate concentrations. Enzyme kinetics at different substrate concentrations are shown in red. Enzyme kinetics at different oxygen concentrations are shown in blue. Kinetic data are extracted from[5,6].

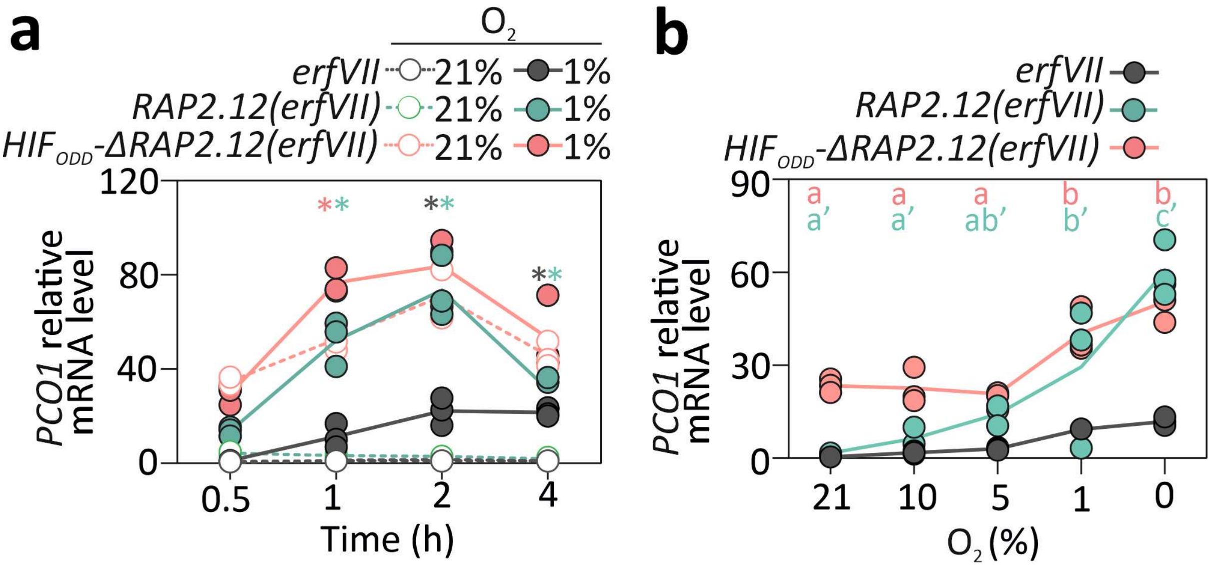


**Figure S10. PCO1 gene expression in the *erfVII* background complemented with the original and chimeric oxygen sensing systems. a,** Expression of the hypoxia marker gene *PCO1* in 7-day-old Arabidopsis seedlings grown under normoxia and then subjected to 21% or 1% $O_2$ for 0.5 to 4 hours. Statistical significance of gene expression relative to controls is denoted by asterisks: grey for *erfVII*, pink for HIF-ΔRAP2.12-2xHA and green for RAP2.12-2xHA. **b,** Dynamics of *PCO1* marker gene expression in response to varying $O_2$ levels in plants expressing RAP2.12 under the control of the PCO or PHD pathways in an *erfVII* background. Letters indicate statistical significance across genotypes, analysed by two-way ANOVA for different $O_2$ percentages, followed by Tukey's post-hoc test for multiple comparisons.

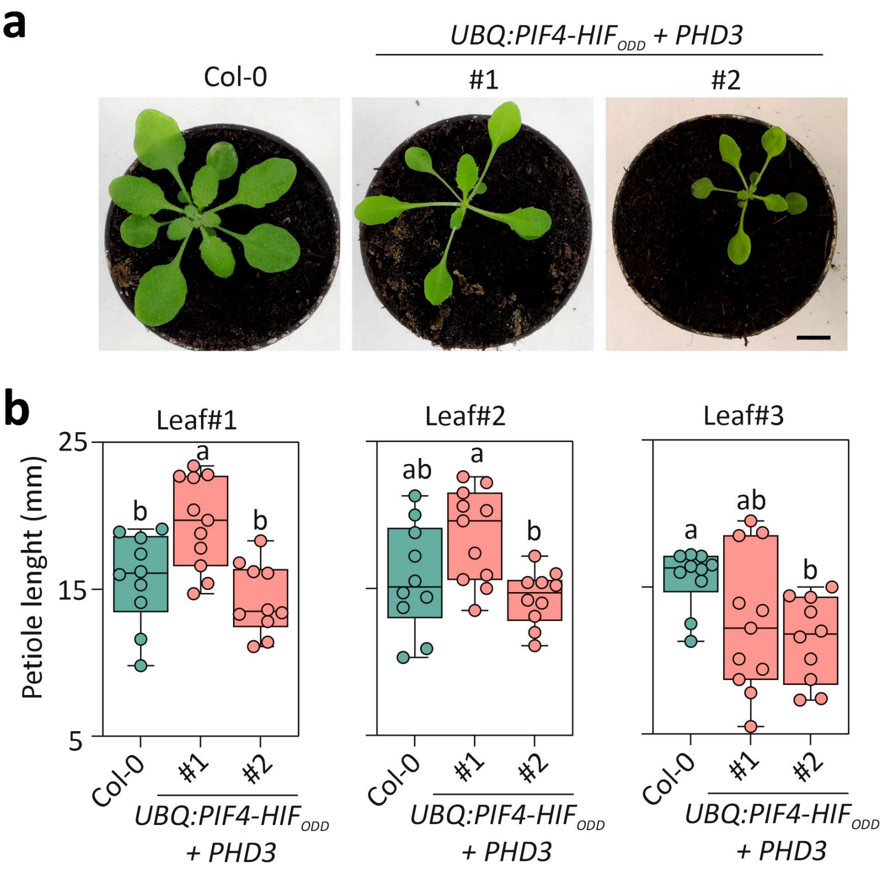


**Figure S11. Characterisation of Arabidopsis expressing PIF4-HIF$_{ODD}$ under control of the *UBQ10* promoter. a,** Phenotypes of wild-type Col-0 plants and two independently transformed plants expressing *PIF4-HIF*$_{ODD}$ under the control of the constitutive UBQ10 promoter (Scale = 1 cm). **b,** Quantification of petiole length of leaves 1 to 3 in *PIF4-HIF*$_{ODD}$ plants. Statistical significance, as determined using two-way ANOVA followed by Tukey's post-test ($p \leq 0.05$) is indicated by letters.

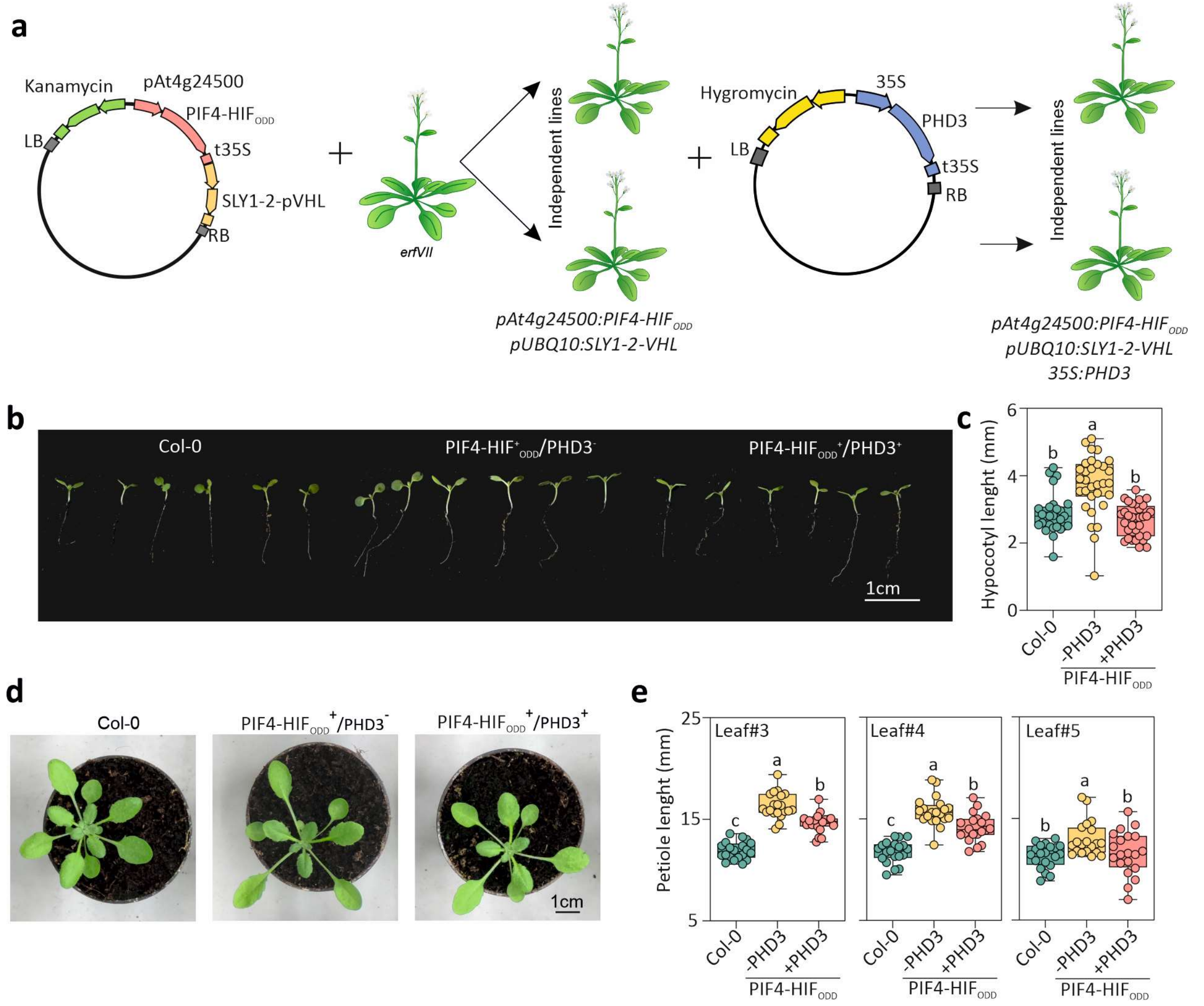


**Figure S12. The PIF4-HIF$_{ODD}$ chimera regulates organ elongation in an $O_2$ dependent manner.** **a,** Implementation of flooding-conditional elongation was achieved in two steps. First, plants were transformed with a construct bearing PIF4-HIF$_{ODD}$ and SLY1-2-VHL. Subsequently, two independent lines were super-transformed with a binary vector containing *35S:PHD3* to control PIF4-HIF$_{ODD}$ stability. **b,** Phenotype of seedlings expressing PIF4-HIF$_{ODD}$ and SLY1-2-VHL in the presence or absence of the PHD3 enzyme. **c,** Hypocotyl length of plants expressing PIF4-HIF$_{ODD}$ and SLY1-2-VHL in the presence or absence of the PHD3 enzyme. Statistical significance, as determined using two-way ANOVA followed by Tukey's post-test ($p \leq 0.05$) is indicated by letters. **d,** Phenotypes of three-week-old Arabidopsis Col-0 and transgenic lines carrying PIF4-HIF$_{ODD}$ with or without PHD3. **e,** Petiole length of plants expressing PIF4-HIF$_{ODD}$ and SLY1-2-VHL in the presence or absence of the PHD3 enzyme. Statistical significance, as determined using two-way ANOVA followed by Tukey's post-test ($p \leq 0.05$) is indicated by letters.

**Supplementary Table S2: List of plasmids used in this study**

| Plasmid | Description |
|---|---|
| 35S:TIR1$_{Fbox}$-VHL (small for protopalsts) | This study |
| 35S:HIF-GAL4-AD (small for protoplasts) | This study |
| 35S:PHD3 | [7] |
| prom4xUAS:FLUC (small for protoplasts) | [8] |
| 35S:SLY1-VHL | This study |
| 35S:SLY1-2-VHL | This study |
| 35S:SLY1-10-VHL | This study |
| 35S:SLY1-2-VHL-GFP | This study |
| 35S:Ω:RLuc-UBQ-HIF$_{ODD}$-Fluc | This study |
| 35S:Ω:RLuc-UBQ-HIF$_{ODD}$-Fluc-nls | This study |
| 35S:Ω:RLuc-UBQ-HIF$_{ODD}$-Fluc-HIF$_{ODD}$ | This study |
| 35S:ADH:RLuc-UBQ-HIF$_{ODD}$-Fluc-HIF$_{ODD}$ | This study |
| 35S:ADH:RLuc-UBQ-HIF$_{ODD}$-Fluc | This study |
| 35S:Ω:FLuc-UBQ-HIF$_{ODD}$nLuc | This study |
| 35S:Ω:FLuc-UBQ-HIF$_{ODD}$nLucHIF$_{ODD}$ | This study |
| 35S:ADH:FLuc-UBQ-HIF$_{ODD}$nLuc | This study |
| 35S:ADH:FLuc-UBQ-HIF$_{ODD}$nLucHIF$_{ODD}$ | This study |
| 35S:PHD3(R205K) | This study |
| 35S:PHD3(D137H) | This study |
| 35S:PHD3(D137E) | This study |
| 35S:PHD3(D315E) | This study |
| 35S:PHD3(R383K) | This study |
| promRAP2.12:HIF$_{ODD}$-2xHA-Δ13RAP2.12/SLY1-2-VHL | This study |
| promRAP2.12RAP2.12-2xHA | This study |
| promPCO4:PHD3 | This study |
| pUBQ10:PIF4-HIF$_{ODD}$ | This study |
| pRON3:PIF4-HIF$_{ODD}$ | This study |

**Supplementary Table S3: List of oligonucleotides used for cloning**

| Name | Sequence |
|---|---|
| 3xHA_Fw | caccatggggcaacaaggatacccg |
| UBQ10_Rv | ctttatgtttttggcgtcttcaccacctcttaacctg |
| FLuc_Fw | caggttaagaggtggtgaagacgccaaaaacataaag |
| FLuc_Hif_Rv | tcaagatctaagttgaaaatcatcatccataggaatataaggagcaag catttcaagatcaagatcagtatccaatttggactttccgcccttctt ggcc |
| p16_Fw | gggaggcctaatggaaccatcttttg |
| p16_Rv | cccctagttttcagagcaggaagg |
| NcoI_pRAP_Fw | gggccatggcatgatggatatgaac |
| NcoI_pRAP_Rv | cccccatggggcggcgattcttgagg |
| ApaI_pOLE_Fw | ccgggcccgtatgtaggtatagtaac |
| CpoI_mCHERRY_Rv | ggggcggtccgtcacttgtgccccag |
| ApaI_At2S3_Fw | cgggcccggaggaaaccaaattaac |
| CpoI_t35S_Rv | ggtccgggtgatctgacgcctcg |
| PHD3Asp137His_Fw | atgttagacatgttcataatcctaatggagatgg |
| PHD3Asp137His_Rv | ccatctccattaggattatgaacatgtctaacat |
| PHD3Asp137Glu_Fw | atgttagacatgttgaaaatcctaatggagatgg |
| PHD3Asp137Glu_Rv | ccatctccattaggattttcaacatgtctaacat |
| PHD3Arg205Lys_Fw | ccttcttatgctactaaatatgctatgactgtt |
| PHD3Arg205Lys_Rv | aacagtcatagcatatttagtagcataagaagg |
| StuI_pAt4g24500_Fw | aggccttcttctccgacgactgtgag |
| SpeI_pAt4g24500_Fw | actagttgtgattcaactacaagatcaaaagg |
| At1G43800_F (SAD6) | ttggcaacccgcttctttcttacc |
| At1G43800_R (SAD6) | tttccctcagctcacgaacctg |
| At5g15120_F (PCO1) | attgggtggttgatgctccaatg |
| At5g15120_R (PCO1) | atgcatgttcccgccatcttc |

**Supplementary Table 4. Reactions and reaction rates of mathematical model**

| | Reactions | Reaction Rates | Description |
|---|---|---|---|
| $v_1$ | → RAP2.12 | $k_1$ | RAP2.12 synthesis |
| $v_2$ | RAP2.12 → null | $k_2 \cdot [RAP2.12]$ | RAP2.12 degradation |
| $v_3$ | RAP2.12 + $O_2$ + PCO4 → PCO4 | $k_3 \cdot [PCO4] \cdot \frac{[O_2]}{Km_1+[O_2]} \cdot \frac{[RAP2.12]}{Km_2+[RAP2.12]}$ | RAP2.12 degradation through oxidation by PCO4 |
| $v_4$ | RAP2.12 → mRNA + RAP2.12 | $k_4 \cdot \frac{[RAP2.12]}{Km_3+[RAP2.12]}$ | HRG mRNA production by RAP2.12 |
| $v_5$ | mRNA → null | $k_5 \cdot [mRNA]$ | HRG mRNA decay |
| $v_6$ | → $HIF_{ODD}$-ΔRAP2.12 | $k_1$ | $HIF_{ODD}$-ΔRAP2.12 synthesis |
| $v_7$ | $HIF_{ODD}$-ΔRAP2.12 → null | $k_2 \cdot [HIF_{ODD}\text{-}\Delta RAP2.12]$ | $HIF_{ODD}$-ΔRAP2.12 degradation |
| $v_8$ | $HIF_{ODD}$-ΔRAP2.12 + $O_2$ + PHD → PHD | $k_3 \cdot [PHD] \cdot \frac{[O_2]}{Km_1+[O_2]} \cdot \frac{[HIF_{ODD}-\Delta RAP2.12]}{Km_2+[HIF_{ODD}-\Delta RAP2.12]}$ | $HIF_{ODD}$-ΔRAP2.12 degradation through hydroxylation by PHD |
| $v_9$ | $HIF_{ODD}$-ΔRAP2.12 → mRNA + $HIF_{ODD}$-ΔRAP2.12 | $k_4 \cdot \frac{[HIF_{ODD}-\Delta RAP2.12]}{Km_3+[HIF_{ODD}-\Delta RAP2.12]}$ | HRG mRNA production by $HIF_{ODD}$-ΔRAP2.12 |

**Supplementary Table 5. Ordinary differential equations of the kinetic model. The reaction rates v1– v9 are given in Supplementary Table S4.**

| | Left-hand Sides | Right-hand Sides | Simulated Initial Concentrations (μM) |
|---|---|---|---|
| RAP2.12/PCO4 model | d[RAP2.12]/dt | v1-v2-v3 | 0.0009 |
| | d[RNA]/dt | v4-v5 | 0.0723 |
| $HIF_{ODD}$-ΔRAP2.12/PHD model | d[$HIF_{ODD}$-ΔRAP2.12]/dt | v6-v7-v8 | 0.0058 |
| | d[RNA]/dt | v9-v5 | 0.4407 |

**Supplementary Table 6. Parameter values used in the kinetic model.** Concentrations and the Michaelis-Menten constants (Kms) are given in μM, except Kms for $O_2$ are in %. Rate constants (ks) are expressed in $s^{-1}$, except $k_1$ is nM $s^{-1}$.

| Parameters | Description | Values | References |
|---|---|---|---|
| $k_1$ | RAP2.12/$HIF_{ODD}$-ΔRAP2.12 protein synthesis rate | 0.005 | [9,10] |
| $k_2$ | RAP2.12/$HIF_{ODD}$-ΔRAP2.12 degradation rate | 0.0002 | [10] |
| $k_3$ | Catalytic rate constant for PCO4-mediated oxidation of RAP2.12 | 26.5878 | [5]Re-fitted |
| $Km_1$ | Michaelis-Menten constant for $O_2$ as a substrate of PCO4 | 16.4217 | [5]Re-fitted |
| $Km_2$ | Michaelis-Menten constant for RAP2.12 as a substrate of PCO4 | 270.2505 | [5]Re-fitted |
| PCO4/PHD | PCO4/PHD protein concentration | 0.1 | [11] |
| $k_4$ | Activation rate constant for HRG expression by RAP2.12 | 0.0016 | [12] |
| $Km_3$ | Michaelis-Menten constant for HRG expression | 0.05 | Assumed |
| $k_5$ | HRG mRNA decay rate | 0.00038 | [10] |
| $k_6$ | Catalytic rate constant for PHD-mediated oxidation of $HIF_{ODD}$-ΔRAP2.12 | 0.1185 | [6]Re-fitted |
| $Km_4$ | Michaelis-Menten constant for $O_2$ as a substrate of PHD | 27.3944 | [6]Re-fitted |
| $Km_5$ | Michaelis-Menten constant for $HIF_{ODD}$-ΔRAP2.12 as a substrate of PHD | 7.8463 | [6]Re-fitted |
